# TRPV4 modulation affects mitochondrial parameters in adipocytes and its inhibition upregulates lipid accumulation

**DOI:** 10.1101/2024.04.21.590448

**Authors:** Shamit Kumar, Tusar Kanta Acharya, Satish Kumar, Young-Tae Chang, Chandan Goswami

## Abstract

Enhanced lipid-droplet formation by adipocytes is a complex process and relevant for obesity. Using knock-out animals, involvement of TRPV4, a thermosensitive ion channel in the obesity has been proposed. However, exact role/s of TRPV4 in adipogenesis and obesity remain unclear and contradictory. Here we used *in vitro* culture of 3T3L-1 preadipocytes and primary murine-mesenchymal stem cells as model systems, and a series of live-cell-imaging to analyse the direct involvement of TRPV4 exclusively at the adipocytes that are free from other complex signalling as expected in *in-vivo* condition. Functional TRPV4 is endogenously expressed in pre- and in mature-adipocytes. Pharmacological inhibition of TRPV4 enhances differentiation of preadipocytes to mature adipocytes, increases expression of adipogenic and lipogenic genes, enhances cholesterol, promotes bigger lipid-droplet formation and reduces the lipid droplet temperature. TRPV4 regulates mitochondrial-temperature, Ca^2+^-load, ATP, superoxides, cardiolipin, membrane potential (ΔΨm), and lipid-mitochondrial contact sites. TRPV4 also regulates the extent of actin fibres, affecting the cells mechanosensing ability. These findings link TRPV4-mediated mitochondrial changes in the context of lipid-droplet formation involved in adipogenesis and confirms the direct involvement of TRPV4 in adipogenesis. These findings may have broad implication in treating adipogenesis and obesity in future.

**Summary Statement:** Thermosensitive ion channel TRPV4 regulates adipogenesis and changes mitochondria metabolism to facilitate bigger lipid droplet formation

## Introduction

TRPV4 is a non-selective thermosensitive cation channel and is endogenously expressed in diverse cells [1]. Notably, TRPV4 has been detected in different stem cells and in cells with different lineages. TRPV4 is known to be a regulator of various physiological processes such as osteoblastogenesis, osteoclastogenesis, and also adipogenesis [2–4]. Presence of TRPV4 is already established in adipose tissues and also in 3T3L-1 pre-adipocytes [5]. Genetic deficiency of TRPV4 (*Trpv4^-/-^* animals) alters the adipogenic differentiation potential of mesenchymal and adipose-derived stem cells [6]. In humans, several TRPV4 variants are shown to be associated with BMI and obesity [7]. However, the involvement of TRPV4 in obesity and in adipogenesis is still not clear. In fact, there are contradictory reports on the role of TRPV4 in the context of adipogenesis. TRPV4 KO mice is shown to be susceptible to high fat diet induced obesity and weight gain [6]. This in general suggest the involvement of TRPV4 in functions that consume energy. Contradictory studies show that TRPV4 KO mice are resistant to high fat diet-induced weight gain [2]. Notably, at least one copy of TRPV4 is retained in all vertebrates [8]. This impose the serious need for further studies to elucidate the exact role of TRPV4 in the context of adipogenesis, especially in adipocytes. This also needs the use of isolated adipose cells *in vitro* condition/s which is free from diverse factors such as adipokines, cytokines, inflammatory factors etc. secreted by other cells.

Mitochondria are the dynamic organelles of cell that are involved in wide plethora of processes such as ATP and ROS production, Ca^2+^-homeostasis, etc. [9]. Mitochondrial regulation and functions are central to brown adipocytes, especially in the context of non-shivering thermogenesis [10]. Mitochondrial functions are also critical for changes triggered during the adipogenesis and also for the overall adipogenesis. Mitochondrial biogenesis is crucial for adipocytic differentiation [11]. Mitochondrial metabolism related parameters such as generation of ATP, ROS, membrane potential, and cardiolipin are known regulator of adipogenesis [12–15]. In spite of substantial research on mitochondria, still little is known about the regulation of mitochondria and subtle changes involved in it during adipogenesis.

Recently, we have reported the presence of TRPV4 in mitochondria where it changes mitochondrial Ca^2+^, temperature and metabolism properties in different cell types [16, 17]. TRPV4 also regulates the ER-mito contact points relevant for cellular (and sub-cellular) Ca^2+^-buffering and Ca^2+^- signalling [17, 18]. Thus we evaluated if and how TRPV4 modulation affects mitochondrial parameters during adipogenesis that may in turn affect the extent of adipogenesis. In this study, we used 3T3L-1 pre-adipocyte cell line and mesenchymal derived primary cells to address the involvement of endogenously expressed TRPV4 in adipogenesis. Our studies indicate that inhibition of TRPV4 function enhances the adipogenesis process and regulates mitochondrial metabolism related properties during it.

## Material and Method

### Reagents and Antibody

KO validated anti-TRPV4 antibody was purchased from Alomone (Cat #: ACC-034, Israel). Monoclonal anti-PPARγ (#C26H12) and anti-SCD1 (#C12H5) antibody were purchased from Cell Signalling Technology. Anti-rabbit secondary antibody coupled to Alexa Fluor488 was purchased from Invitrogen. Insulin, IBMX, and Rosiglitazone were purchased from Merck (Sigma Aldrich). Oil Red O powder was obtained from HiMedia. Mito Thermo Yellow (MTY), Droplet Thermo Green (DTG) and ER Thermo Yellow (ETY) were synthesized as described previously [19–21]. Fluo-4-AM was purchased from Thermo Fisher Scientific.

### Cell culture, transfection and pharmacological modulation of TRPV4

3T3L-1 cells were procured from NCCS (Pune, India). Cells were grown in RPMI 1640 GlutaMAX supplement media (GIBCO) containing FBS (10% v/v), Streptomycin (100μg/ml), and Penicillin (100U/ml) in humidity-controlled incubator maintained at 5% CO_2_ and 37°C. For transient transfection, Lipofectamine 3000 Plus reagent (Invitrogen) was used according to manufactures protocol. Confocal imaging was performed 24 hours post transfection. For modulation of TRPV4, 4αPDD (agonist) and RN1734 (antagonist) was used at a concentration of 5μM and 10μM respectively. Subsequently, cells were either washed for live cell imaging or fixed with 4% PFA.

### Isolation and culture of primary murine mesenchymal stem cells (MSC’s)

Isolation of primary bone marrow cells was performed as previously reported [22]. This study was approved by the Institutional Animal Ethics Committee (IAEC) (NISER/SBS/IAEC/AH-55). Briefly, adult male BALB/c mice (4-6 weeks old) were sacrificed by CO_2_ asphyxiation. Bone marrow from tibia and femur was flushed out with culture media (RPMI 1640 with 10% FBS) and cells were passed through 70μm cell strainer (SPL Labs). Subsequently, RBC lysis was performed and cells were plated in 100 mm dish (Thermo Fisher Scientific) and grown in CO_2_ incubator. After 1 day of incubation, the non-adherent hematopoietic cells are removed and the adherent cells are grown till confluence. After 1-2 passages, majority of the cells were mesenchymal stem cells as identified by their fibroblast-like morphology and used for the study.

### Immunocytochemistry, live cell imaging and microscopy

For immunocytochemistry, cells were seeded on 18mm coverslips. Either differentiation of pre-adipocyte was carried out or after 50% confluency, 4αPDD and RN1734 was added for 24 hours. Cells were fixed with 4%PFA at room temperature. Cells were subsequently washed with PBS thrice and cell permeabilization was performed using TritonX-100 (0.1%) in PBS for 5 minutes followed by washing with PBS twice. Blocking was done by using 5% Bovine Serum Albumin (BSA) in PBS for 1 hour. Subsequently, cells were washed and incubated with primary antibody (against TRPV4, PPARγ, or SCD1 at dilution of 1:500 for all) and kept overnight at 4° C. Following this, cells were washed thrice with PBS and further incubated with secondary antibody (anti-rabbit) at 1:500 dilutions along with Phalloidin-594 at 1:500 dilution in PBS and 5% BSA at 1:1 ratio for 2 hours. Following washing, cells were incubated with DAPI (5μM) in PBS for 10 minutes at room temperature and coverslips were then washed and mounted on glass slides (Globe Scientific) using Fluoromount G (Southern Biotech). For live cell imaging, cells were seeded on 25mm coverslips and grown till 50% confluency before treating them with modulators of TRPV4. Cells were imaged using confocal microscope (FV3000, Olympus) with its 63X objective. For comparative analysis, all the images were taken using same parameters. ImageJ was employed for image analysis and quantification. In each case the intensity/area was calculated (unless stated otherwise) using the ‘Measure’ function. The images are displayed in intensity profile (Rainbow RGB) scale. In case of PPARγ, the nuclear intensity/area was quantified by taking ROI’s from DAPI.

### Adipocyte differentiation

3T3L-1 differentiation was performed by method as described previously [23]. Briefly, 3T3L-1 pre-adipocyte cells were seeded on either 18 mm or 25 mm coverslips in complete media containing 10% (v/v) FBS. Cells were grown till 90-95% confluence before differentiation media was added. Adipogenic differentiation was carried out using differentiation media containing complete media with IBMX (0.5mM), dexamethasone (0.25μM), rosiglitazone (2μM) and insulin (1 μg/ml). Cells were kept in differentiation media with or without TRPV4 modulators for 48 hours. We refer to these as the “Induced Adipocytes”. After 48 hours, maintenance media containing complete media with insulin (1 μg/ml) with or without modulators of TRPV4 was added. Subsequently, media was refreshed after 48 hours until intracellular lipids were visible after differentiation. We refer these cells as the “Mature Adipocytes”. The primary mesenchymal stem cells were seeded on 35 mm glass bottom dishes (IBIDI) and grown till optimum confluence is achieved. For differentiation, complete media containing IBMX (0.5 mM), insulin (10 μg/ml), rosiglitazone (2 μM) and dexamethasone (1 μM) was used for 3 days with or without modulator of TRPV4. Following this, cells were kept in complete media containing insulin (10 μg/ml) for 3-7 days when lipid droplets were visible.

### Oil Red O staining and cholesterol quantification

Oil Red O (ORO) staining was performed as described previously (Yang et al., 2011). Briefly, 3T3L-1 pre-adipocytes were seeded on 35 mm glass bottom dishes (IBIDI) and grown till 90-95% confluence. Adipocyte differentiation was carried out and mature adipocytes were fixed using 4% PFA at room temperature after washing with 1XPBS twice. ORO-stock solution was prepared containing 0.3% (w/v) ORO (HiMedia) in 100% isopropanol. ORO working solution was prepared freshly each time by adding stock solution and MQ water at 3:2 ratio and then filtered using 0.22 μm syringe filter. PFA was aspirated out and cells were washed with 1XPBS thrice and incubated with 60% isopropanol for 5 minutes. ORO working solution was added and incubated for 30 minutes and subsequently washed with 60% isopropanol to eliminate excess ORO stains. Imaging was performed by using confocal microscope with far-red laser. For the quantitative analysis of cholesterol, we used enzyme assay based ‘Cholesterol Quantitation Kit-MAK043’ from Sigma-Aldrich as per the manufacturer’s instructions. The total concentration of the free cholesterol was determined by taking absorbance at 570nm.

### Labelling of cells with thermo-sensing and other dyes

3T3L-1 pre-adipocytes were seeded in either 25mm coverslips or glass bottom dishes (IBIDI) and grown till 50% confluence or differentiated into adipocytes with TRPV4 modulators. In case of pre-adipocytes, 24 hours drug treatment was given and then imaging was done. For induced adipocytes, 3T3L-1 cells were kept in differentiation media for 48 hours and then labelled with different mitochondrial dyes whereas for mature adipocytes, cells were subsequently placed in maintenance media for 48-96 hours and then imaged. TMRM (Tetramethylrhodamine, Methyl Ester, Percholrate) dye at 50 μM concentration for 30 minutes was used for mitochondrial membrane potential analysis. Dye loaded cells were washed and imaged. MitoSOX was used at a concentration of 2.5μM for 15 mins and cells were imaged for comparing mitochondrial superoxide levels. NAO (Nonyl Acridine Orange) dye was used at a concentration of 0.5 μM for 15 mins for checking mitochondrial cardiolipin levels. MitoTrackerRed-CMXRos was used to at 0.5μM concentration for 15 mins to label the mitochondria for morphological analysis. Mito morphology plugin was used in ImageJ to quantify the mitochondria morphological parameters. Mitochondrial ATP levels were measured using the ATP-Red dye at a concentration of 5μM for 15 mins. For measuring the temperature of mitochondria, ER and lipid droplets; Mito-Thermo-Yellow (MTY), ER-Thermo-Yellow (ETY) and Droplet-Thermo-Green (DTG) dye were used respectively, both at concentration of 0.5 μM for 15 mins. In each of the above condition, the images are represented as intensity profile in rainbow RGB scale and intensity/area was quantified for quantitative comparison.

### Live cell lipid staining and quantification of lipid-mitochondrial contact points

Live cell staining of lipid droplet was performed by using BODIPY493/503 (4,4-Difluoro-1,3,5,7,8-Pentamethyl-4-Bora-3a,4a-Diaza-*s*-Indacene) at concentration of 1.9 μM for 30 minutes. Primary murine MSC’s were seeded on 35 mm glass bottom dishes and adipogenic differentiation was carried out. For live cell intracellular lipid staining, 2X solution of BODIPY (3.8 μM) was prepared in complete media and vortexed and used at 1:1 dilution with media. In case of lipid-mito contact points analysis, the labelling of the mitochondria was performed by MitoTrackerRed-CMXRos (0.5μM; 15mins). Subsequently, live cell confocal imaging was performed. For quantification of lipid-mitochondrial contact points, the particle-based quantification approach was considered as previously described [18]. In brief, the colocalization of lipid and mitochondria was done using the ‘Co-localisation’ plugin in Image J. The images were converted into binary format and ‘Analyse Particle’ function was used to quantify the lipid, mitochondria and colocalised particles. For each case, the particle numbers are plotted using GraphPad Prism 8.

### Cellular and mitochondrial Ca^2+^-imaging

For intracellular Ca^2+^ imaging, GCaMP construct was used. GCaMP increases intensity with an increase in the cellular Ca^2+^ level. Cells were transfected with GCaMP and using confocal microscope, live cell time series imaging was done 24 hours after transfection. Agonist and antagonist were added at the 20^th^ frame of the time series imaging. For mitochondrial Ca^2+^-imaging, MitoPericam construct was used (A kind gift from Dr. Atsushi Miyawaki, Saitama, Japan) which enters the mitochondria and upon binding to Ca^2+^ chances its excitation from 415nm to 495nm whereas emission remains at 515nm [24]. MitoPericam increases its intensity with increase in Ca^2+^-levels of mitochondria. MitoPericam construct was transfected into 3T3L-1 pre-adipocytes and time series imaging was done. In case of RN1734 pre-treated condition, RN1734 (10 μM) was added for 1 hour and then imaging was done. For measuring basal mitochondrial Ca^2+^-levels, post transfection modulators of TRPV4 were added for 24 hours and live cell imaging was done. Fluo-4AM dye was used for detecting cellular Ca^2+^-level after treating the cells with 4αPDD and RN1734 for 24 hours. Dye was incubated for 30 minutes at 37°C and subsequently washed with 1xPBS and taken for live cell imaging. To quantify the mitochondrial Ca^2+^-levels, ‘Analyse particle’ function of Image J was used to quantify the intensity and particle numbers. For GCaMP, intensity of 200 frames was quantified using Image J ‘Multi-measure’ function.

### Statistical tests

GraphPad Prism 8 was used for statistical & correlation analysis and plotting of graphs. Data shown are mean ± SEM. One-way ANOVA was performed for statistical comparisons.

## Results

### Differential expression of TRPV4 and organization of F-actin during adipogenesis

We detected endogenous expression of TRPV4 in 3T3L-1 pre-adipocytes (**Fig 1a&d**). The TRPV4 specificity was confirmed using the TRPV4 specific peptide block (**Fig S1a**). Further, we treated the cells with TRPV4-agonist (4αPDD, 5μM) and antagonist (RN1734, 10μM) for 24 hours. No significant differences in the expression of TRPV4 were observed due to pharmacological modulation of TRPV4. We investigated if the expression of TRPV4 changes during the course of adipogenesis and/or if activation or inhibition of TRPV4 affects it’s expression. For this we analysed two time points during adipogenesis. We differentiated 3T3L-1 in presence of 4αPDD and RN1734 using the differentiation media (2 days) which committed 3T3L-1 to adipogenic lineage [23]. We term this as “Induced Adipocytes” wherein cells are committed to be an adipocyte and on the 5^th^ day of differentiation the parameters are measured [23]. In case of induced adipocytes (**Fig 1b&d**), TRPV4 expression is reduced (0.62 fold) as compared to the pre-adipocytes. Also, activation or inhibition of TRPV4 in induced adipocytes show no changes in expression levels, albeit a non-significant increase in the expression is seen in case of 4αPDD treatment. For “Mature Adipocytes” (MA) (**Fig 1c&d**), the “differentiation media” is changed to “maintenance media” along with modulators of TRPV4 for 2 days until intra-cellular lipid droplets become visible. On the 7^th^ day, parameters were measured for MAs. Basal expression of TRPV4 in MA is upregulated (∼2.04 fold) as compared to day 1. Inhibition by RN1734 during the whole process leads to an enhanced expression of TRPV4 whereas, 4αPDD shows no significant change in the expression patterns. Overall, our data suggests that the expression of TRPV4 decreases and then increases during the course of adipogenesis and TRPV4 inhibition results in more TRPV4 expression in MA.

**Figure 1:**
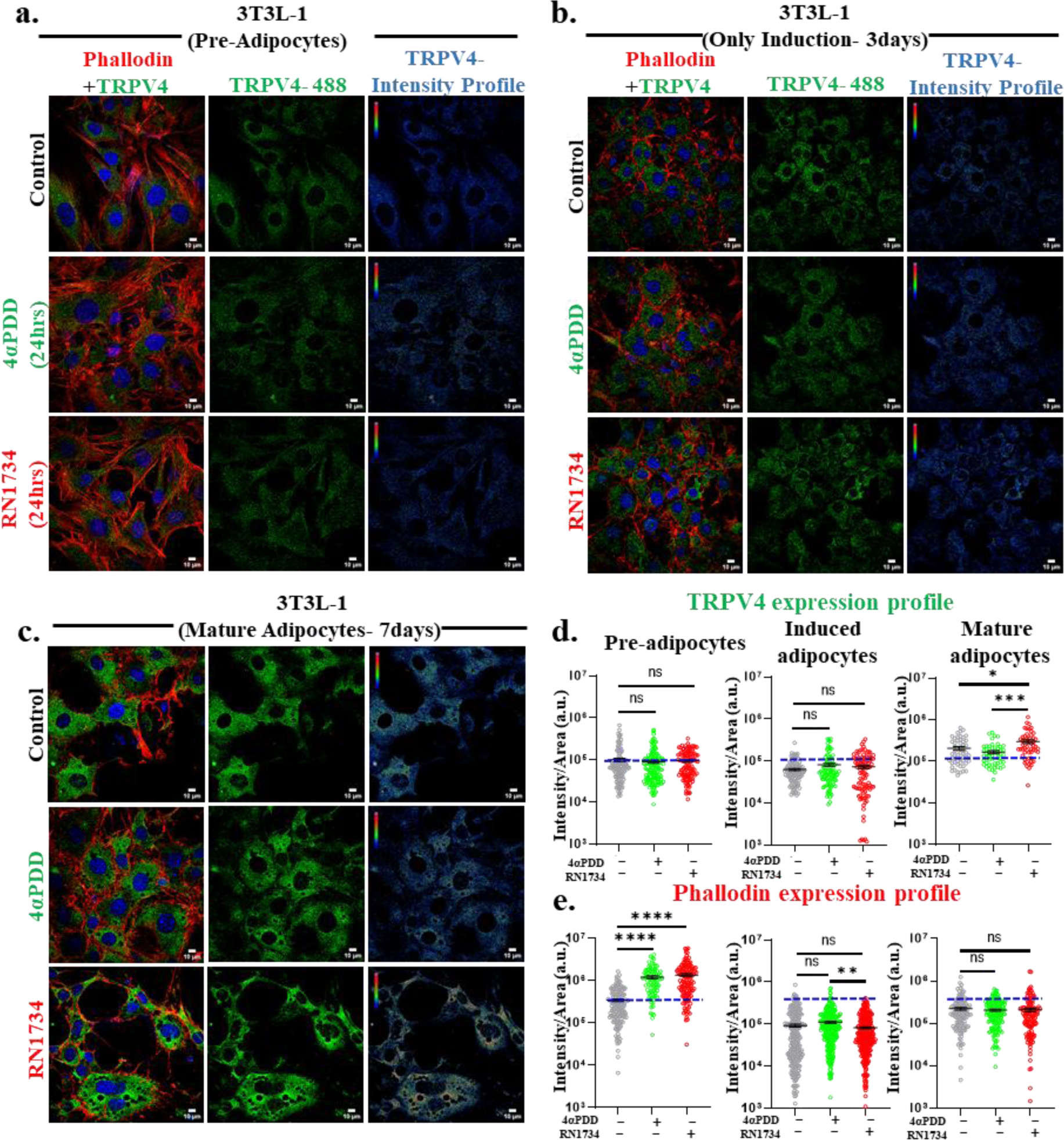
Differential expression of TRPV4 and F-actin during adipogenesis in 3T3L-1. **A.** TRPV4 is endogenously expressed in 3T3L-1 pre-adipocytes. 3T3L-1 pre-adipocytes were treated with TRPV4 agonist 4αPDD (5μM) and antagonist RN1734 (10μM) for 24 hours and stained for TRPV4 (green), F-actin (red), and DAPI (blue). **B.** 3T3L-1 pre-adipocytes were differentiated for 2 days in induction media with or without TRPV4-specific modulators. **C.** 3T3L-1 were differentiated to mature adipocytes for 7 days and expression of TRPV4 was detected using immunofluorescence. **D**. Quantitative analysis of cell-wise TRPV4 expression in pre-adipocyte, induced adipocyte and in mature adipocyte are shown. TPRV4 expression changes during adipogenesis and inhibition of TRPV4 function by RN1734 increases the level of TRPV4 in mature adipocytes. **E.** The F-actin level changes during adipogenesis. Overall, the F-actin level reduces during adipogenesis. n≥50 cells in each condition. One-way ANOVA ****p<0.0001; ***p<0.001; **p<0.01; *p<0.05; ns: non-significant. Scale bar 10μm.

TRPV4 interacts with actin cytoskeleton and also regulates F-actin status causing remodelling of cell morphology [25]. Integrity and nature of the actin cytoskeleton is known to affect adipogenesis [26, 27]. We stained the cells with Phalloidin to evaluate the levels and organization of F-actin (**Fig S1b**). Both activation and inhibition of TRPV4 in pre-adipocytes increase F-actin levels (**Fig 1e**). Further, F-actin level reduces (0.26 fold) in induced adipocytes and does not changes with TRPV4 modulation (**Fig 1e**). However, the F-actin increases in mature adipocytes but still remains less (0.63 fold) than pre-adipocytes. TRPV4 modulation also doesn’t affect the F-actin status of mature adipocytes. Overall, data suggests that during the course of adipogenesis, F-actin level is reduced and TRPV4 modulation does not affect that greatly.

TRPV4 is known to affect the cellular morphology [25]. We checked cell morphological parameters such as area, length and breadth (**Fig S2a**). In pre-adipocytes, both 4αPDD and RN1734 increases the area, length and breadth. In induced adipocytes, all these three parameters decrease. RN1734 reduces area and breadth further in the induced adipocytes. In mature adipocytes, the area, length and breadth increases and are comparable to pre-adipocytes. The area of mature adipocytes is marginally more than pre-adipocytes. TRPV4 modulation does not change the cellular morphology in mature adipocytes. Our data suggests that TRPV4 regulates cell morphology of induced adipocytes, but most-likely act downstream of the actin remodelling.

### Long-term inhibition of TRPV4 leads to elevated cellular Ca^2+^-levels in pre-adipocytes

To check the functionality of TRPV4, we performed Ca^2+^-imaging by using Ca^2+^-sensitive dye Fluo-4-AM. 3T3L-1 pre-adipocytes were treated with either 4αPDD or RN1734 for 24 hours and then measured for cellular Ca^2+^-levels (**Fig 2a**). 4αPDD shows no change on the cellular Ca^2+^-levels, whereas RN1734 increases (∼1.5 fold) the Fluo-4 intensity. In case of 4αPDD, the intensity decreases slightly (0.91 fold) (**Fig S2b**). Thus, long-term inhibition of TRPV4 by RN1734 causes enhanced cytosolic Ca^2+^-level in pre-adipocytes probably due to blockade of intracellular Ca^2+^-buffering (such as by mitochondria and ER).

**Figure 2:**
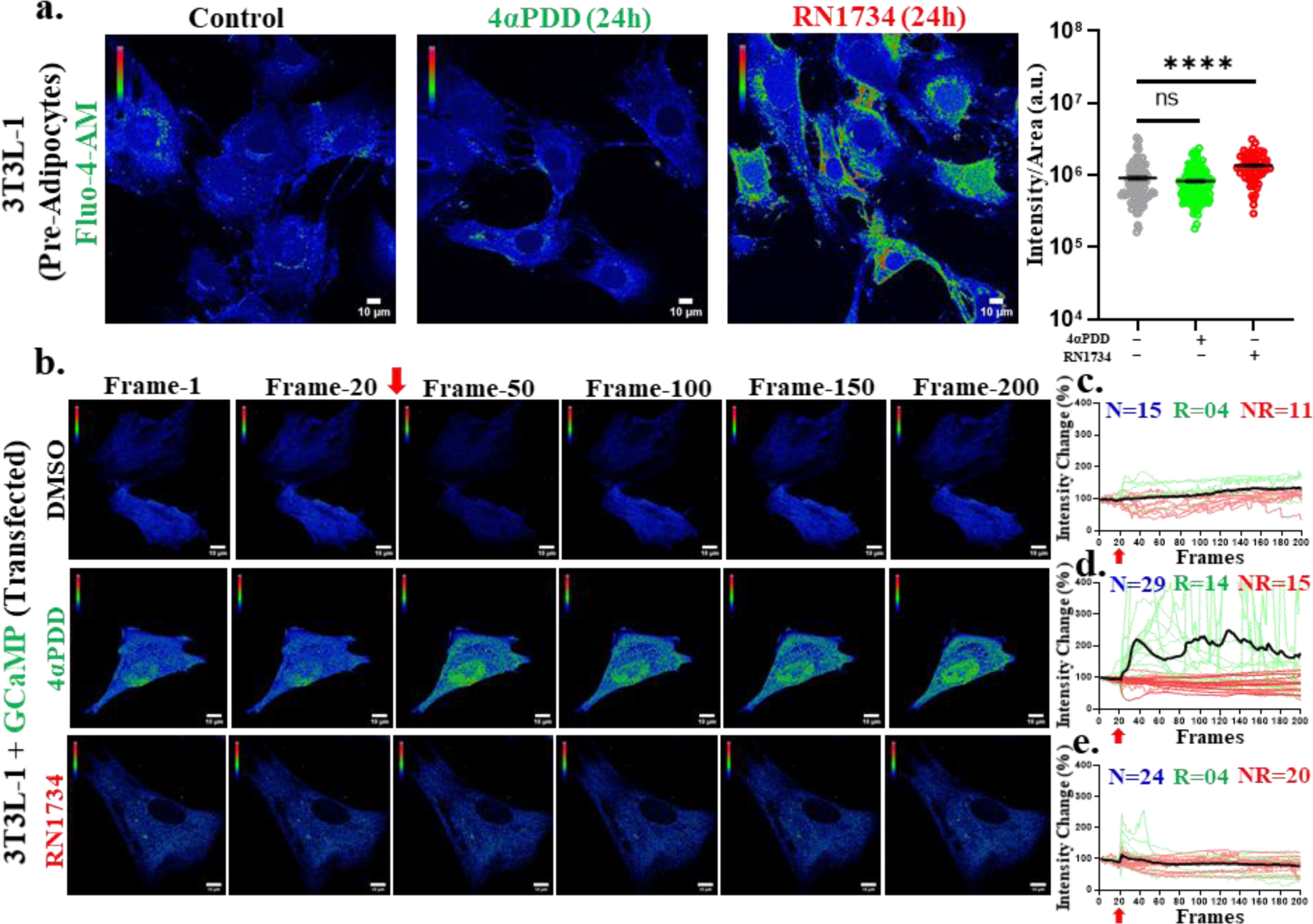
TRPV4 modulation affects Ca^2+^-levels in 3T3L-1 pre-adipocytes. **A.** 3T3L-1 pre-adipocytes were treated with 4αPDD (5μM) and RN1734 (10μM) for 24 hours and labelled with Flou-4-AM dye to measure the level of cellular Ca^2+^. Shown are the intensity profile images in rainbow scale. Long-term inhibition of TRPV4 (by RN1734) increases the cytosolic Ca^2+^ of 3T3L-1 pre-adipocytes. (n>60 cells in each condition). One-way ANOVA ****p<0.0001. Scale bar 10μm. **B**. 3T3L-1 pre-adipocytes were transfected with GCaMP to measure the effect of instantaneous TRPV4 modulation on cytosolic Ca^2+^ levels. Live cell Ca^2+^ imaging was performed for 200 frames (∼3.5 mins) and stimulus (indicated by red arrow) was added at 20^th^ frame of imaging. Shown are the intensity profile images in rainbow scale. Black line represents the average value of all the cells. **C**. Addition of DMSO shows no significant changes in cellular Ca^2+^ level. **D.** 4αPDD (5μM) causes an instantaneous influx of Ca^2+^. **E.** RN1734 (10μM) addition lead to no major alteration in the cellular Ca^2+^ pattern, although slight reduction is seen.

### TRPV4 activation causes an instantaneous Ca^2+^-influx of in pre-adipocytes

We analysed the instant effect of TRPV4 modulation on Ca^2+^-levels and thus performed live cell Ca^2+^-imaging using a genetically encoded Ca^2+^ sensor (GCaMP) (**Fig 2b**). Addition of DMSO shows no significant changes in the cellular Ca^2+^-levels of the 3T3L-1 cells (**Fig 2c**). Addition of 4αPDD causes instant increase in the cellular Ca^2+^ levels which sustains till the end of the experiment (**Fig 2d**). Further, addition of RN1734 shows no notable changes in the Ca^2+^ levels (**Fig 2e**). Upon comparing the normalised change in Ca^2+^-level (200^th^-01^st^ of the average value), we found that 4αPDD addition causes more Ca^2+^-influx (∼2 fold) and RN1734 reduces the cellular Ca^2+^ instantaneously (**Fig S2c**). We classified the cells as responding (R, green) and non-responding (NR, red) on the basis of their changes in the Ca^2+^-levels. We considered a cell as “responding” if it increases its basal Ca^2+^ level by 1.5-fold (>150 intensity change value) after adding stimulus. We found that, the number of responding cells increases in case of 4αPDD (∼1:1 = responding: non-responding cells). Detailed analysis indicates that the entire population contains two distinct groups of cells; one group that contains cells which hyper-responsive to 4αPDD stimulation and other that does not get affected at all by 4αPDD (**Fig S2d**). Altogether, results suggest that activation of TRPV4 causes an instant increase in Ca^2+^, but only in a particular subset of cells.

### TRPV4 inhibition increases differentiation of pre-adipocytes to mature adipocytes (adipocyte turnover)

We evaluated if modulation of TRPV4 affects the extent of adipogenesis. 3T3L-1 pre-adipocytes were differentiated in presence of 4αPDD or RN1734 and ORO-staining was performed to evaluate adipogenesis (**Fig 3a**). The cells that are positive for ORO is considered as truly mature adipocytes (highlighted with yellow arrows). We quantified the % of cells that are truly adipocytes as compared to DAPI positive cells (highlighted by white arrows) when grown under different conditions (**Fig 3a**). We noted that pre-adipocytes when differentiated in control condition (with inducing media), the 74.66% of cells become truly mature adipocytes in 7 days. However, when 4αPDD is present in this process (for last 5 days), only 41.78% cells become ORO-positive. In contrast, presence of RN1734 (for last 5 days) increase the number of cells that have ORO labelling (87.23%). This suggests that inhibition of TRPV4 enhances the differentiation of pre-adipocytes to mature adipocytes or in other way, increases the adipocyte turnover number.

**Figure 3:**
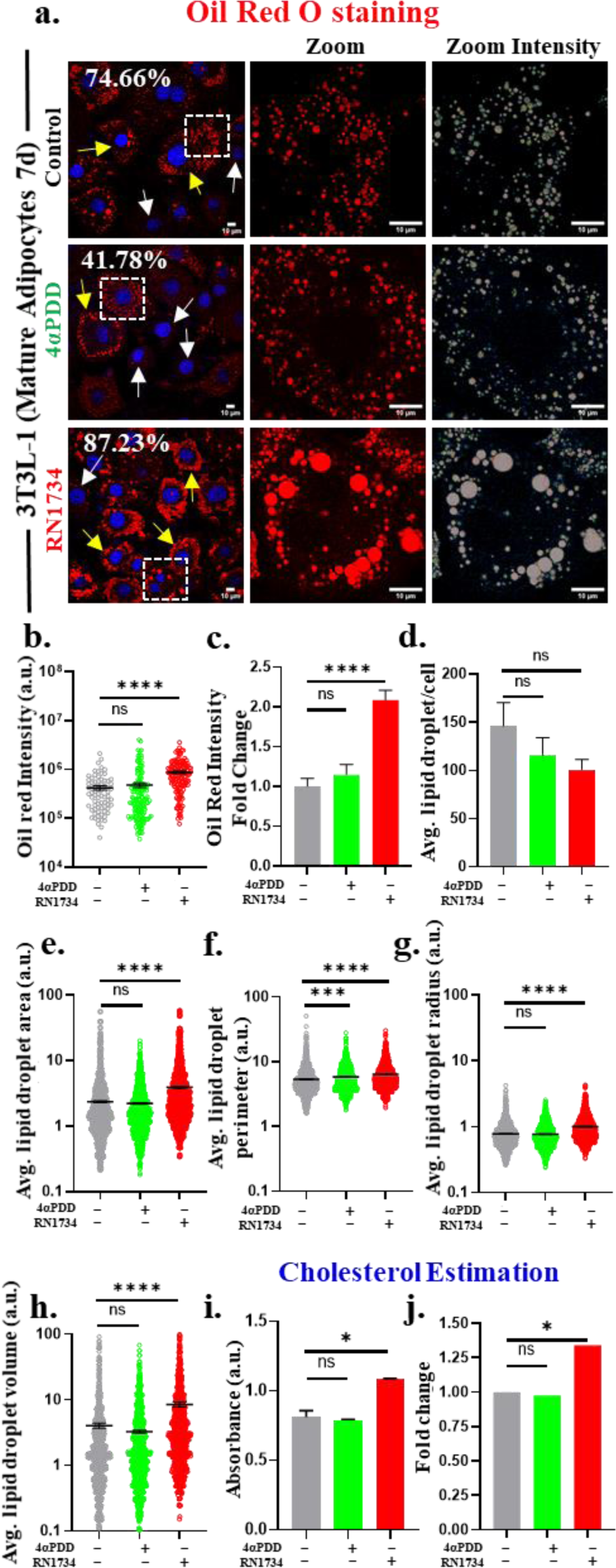
TRPV4 modulation affects the lipid accumulation and cholesterol levels during adipogenesis. **A.** 3T3L-1 were differentiated along with 4αPDD (5μM) and RN1734 (10μM) for 7 days and Oil Red O staining was performed to measure the lipid content of cells. Images were taken using confocal microscope and Oil Red intensity was quantified for each case (n>100 cells for each condition). Number depicts the percentage of Oil Red O positive cells as compared to DAPI positive cells. White arrows depict only DAPI positive but Oil Red O negative cells and yellow arrows depict the Oil Red O positive cells. **B.** TRPV4 inhibition (RN1734) during adipogenesis leads to more lipid content in the cells. **C.** Lipid amount in the cells treated with RN1734 doubles (∼2-fold increase). **D-H.** Individual lipid droplets from different cells was measured for area, number, perimeter, radius, and volume. RN1734 treatment doesn’t affect the lipid droplet number rather increases the size of the lipid droplet as seen by increase in the area, perimeter, radius and volume of lipid droplets. n>1000 lipid droplets were measured in each condition. **I-J.** TRPV4 inhibition during adipogenesis also raises the cholesterol content in adipocytes. A ∼1.25-fold increase in the cholesterol level of cells was seen in RN1734 condition. One-way ANOVA ****p<0.0001; ***p<0.001; *p<0.05; ns: non-significant. Scale bar 10μm.

### TRPV4 inhibition during adipogenesis increases both number and sizes of the lipid droplets

We further quantified the ORO intensity of the different conditions. Presence of 4αPDD during adipogenesis shows no significant changes in the ORO-intensity as compared to control condition (**Fig 3b**). But, in presence of RN1734 (TRPV4 inhibition), the ORO-intensity increases significantly than the control condition which suggests increased adipogenesis (**Fig 3b**). We found that the ORO-intensity increases by 2 fold in RN1734-treated cells suggesting enhanced adipogenesis due to TRPV4 inhibition (**Fig 3c**).

We quantified more than >1000 individual lipid droplets in each condition. We found that there is no significant difference in the average number of lipid droplets per cell although a slight reduction occurs in both 4αPDD and RN1734-treated condition (**Fig 3d**). The average area of lipid droplet goes up significantly in RN1734-treated condition than the control group, but no change is seen in case of 4αPDD-treated condition (**Fig 3e**). In addition, the average perimeter, radius and volume of the lipid droplet increases in RN1734-treated condition (**Fig 3f-h**). No change is seen in these parameters in case of 4αPDD (except the perimeter of lipid droplet; which increases) (**Fig 3f**).

We subsequently plotted the individual lipid droplets in an ascending order of their area. We observed a bi-phasic curve in each condition (**Fig S3a**). However, the size of the “smallest lipid droplet” in RN1734-treated condition is greater than the control and the 4αPDD -treated condition. We subsequently classified the lipid droplets on the basis of their size (area) as small (0-1 a.u., light blue); medium (1-10 a.u., dark blue); large (10-20 a.u., light red); and very large (>20 a.u., dark red) (**Fig S3c**). We observed that the percentage of small lipid droplets decreases greatly in RN1734-treated condition (from 28.77% to 11.40%) and remains almost unchanged in 4αPDD-treated condition (28.18%). This remain the case for lipid droplets with medium area where these droplets increase in RN1734-treated condition (81.37%) from control (68.70%) and 4αPDD (70.44%). The lipid droplets with large (5.15%) and very large (2.08%) area are also increased in RN1734-treated condition as compared to control. In 4αPDD-treated condition, the large and very large lipid droplet decreases (from 1.92% to 1.30% and 0.62% to 0.09% respectively) as compared to control condition. Therefore, TRPV4 modulation during adipogenesis bring about changes in lipid droplet size and its distribution.

We also measured the level of cholesterol in the mature adipocytes. Addition of 4αPDD during adipogenesis does not change the level of cholesterol significantly, but RN1734 leads to a significant rise in the cholesterol levels (**Fig 3i**). We found ∼1.3 fold more cholesterol in RN1734-treated condition (**Fig 3j**). All in all, our data suggests that TRPV4 inhibition during the course of adipogenesis results in enhanced extent of adipogenesis, i.e. increase in size and volume of the lipid droplets and increased cholesterol levels.

### TRPV4 inhibition enhances the expression of adipogenic and lipogenic genes

Since, inhibition of TRPV4 is increasing the adipocyte turnover number and promoting the formation of bigger lipid droplets, we checked for the expression of standard lipogenic marker, namely SCD1 in the induced and mature adipocytes (**Fig 4a**). Inhibition of TRPV4 in induced adipocytes lead to significantly higher level of SCD1 whereas activation of TRPV4 does not affect the SCD1 expression (**Fig 4b**). We noted ∼4-fold increase in the intensity in RN1734-treated cells (**Fig 4c**). In mature adipocytes, the basal level of SCD1 increases as compared to induced adipocytes (∼4.38-folds) (**Fig 4d**). Here also, TRPV4-agonist treated cells have no such changes in the SCD1 expression however, TRPV4-antagonist treated cells shows a significantly increased level of SCD1 (**Fig 4d**). Notably, ∼1.84-fold more intensity is observed in the RN1734-treated cells (**Fig 4e**).

**Figure 4:**
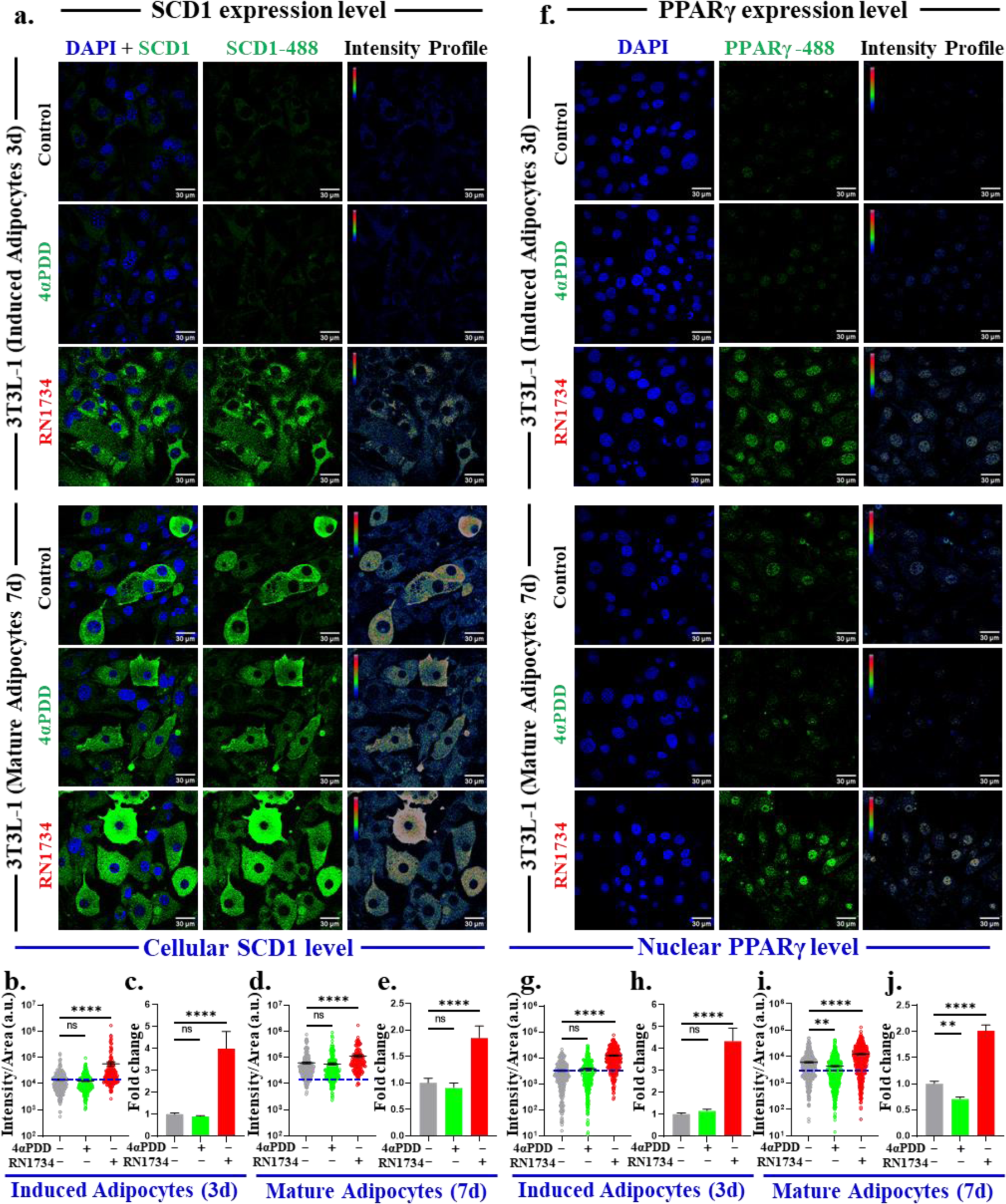
TRPV4 inhibition increases the level of adipogenic and lipogenic marker genes. 3T3L-1 cells were differentiated in presence of 4αPDD and RN1734 and stained for adipogenic marker genes SCD1 and PPARγ. **A.** Shown are the immune-fluorescence images. Cells were compared for level of SCD1 in induced and mature adipocytes. **B-E.** TRPV4 inhibition increases the SCD1 level in both induced adipocytes (∼4-fold) and mature adipocytes (∼1.84-fold). **F.** The levels of PPARγ is also evaluated in these conditions. **G-J.** TRPV4 antagonist treated cells display an elevated level of PPARγ in both induced (∼4-fold rise in intensity) and mature adipocyte cells (∼2-fold rise in intensity). n>250 cells in each of the condition. One-way ANOVA ** = p<0.01; **** = p<0.0001; ns: non-significant. Scale bar 30μm.

In a similar manner, we also evaluated the expression of another adipogenic marker, namely PPARγ (**Fig 4f**). We quantified the amount of PPARγ present in the nucleus in different conditions by taking ROI’s from DAPI. In induced adipocytes, activation of TRPV4 does not affect the nuclear PPARγ level (**Fig 4g**). However, inhibition of TRPV4 increases the amount of nuclear PPARγ significantly (∼4.33-folds increase in the intensity) (**Fig 4g&h**). In mature adipocytes, the basal expression of PPARγ elevates (∼1.88-fold) as compared to induced adipocytes. Here, activation of TRPV4 reduces the level of nuclear PPARγ whereas, inhibition of TRPV4 increases the nuclear PPARγ significantly (**Fig 4i**). A reduction of ∼0.70-fold in intensity is observed in 4αPDD-treated cells and ∼2-fold increase in intensity is observed in RN1734-treated cells (**Fig 4j**). Overall, the data suggests that inhibition of TRPV4 during adipogenesis increases the expression of adipogenic genes. Moreover, the expression of the adipogenic and lipogenic markers drastically increases in TRPV4 inhibitor treated induced adipocytes suggesting more commitment of these cells to adipogenic lineage.

### TRPV4 modulation affects mitochondrial morphometric parameters

We used MitoTrackerRed-CMXRos dye in 3T3L-1 pre-adipocytes and quantified several mitochondrial morphological parameters (**Fig 5a**). Modulation of TRPV4 by 4αPDD or RN1734 (for 24 hours) does not affect the mitochondrial counts or numbers drastically (**Fig 5b**). But, mitochondrial content is increased due to activation of TRPV4, but remain unaffected in case of RN1734 (**Fig 5c**). Also, the average mitochondrial area and perimeter enhanced due to 4αPDD-treatment (**Fig 5e&f**). However, both mitochondrial area and perimeter remain unchanged in case of RN1734 treatment (**Fig 5e&f**). This increment in mitochondrial area and perimeter also correlates with the increase in mitochondrial interconnectivity seen in case of 4αPDD condition (**Fig 5h**). The mitochondria interconnectivity also increases in RN1734-treated cells (**Fig 5h**). No significant change is seen in mitochondrial solidity due to 4αPDD-treatment whereas, RN1734-treatment leads to a significant increase in the mitochondrial solidity (**Fig 5d**). The mitochondrial circularity is not affected by modulation of TRPV4 (**Fig 5g**). Overall results suggest that TRPV4 modulation affects the mitochondrial content and morphology in 3T3L-1 pre-adipocyte cells.

**Figure 5:**
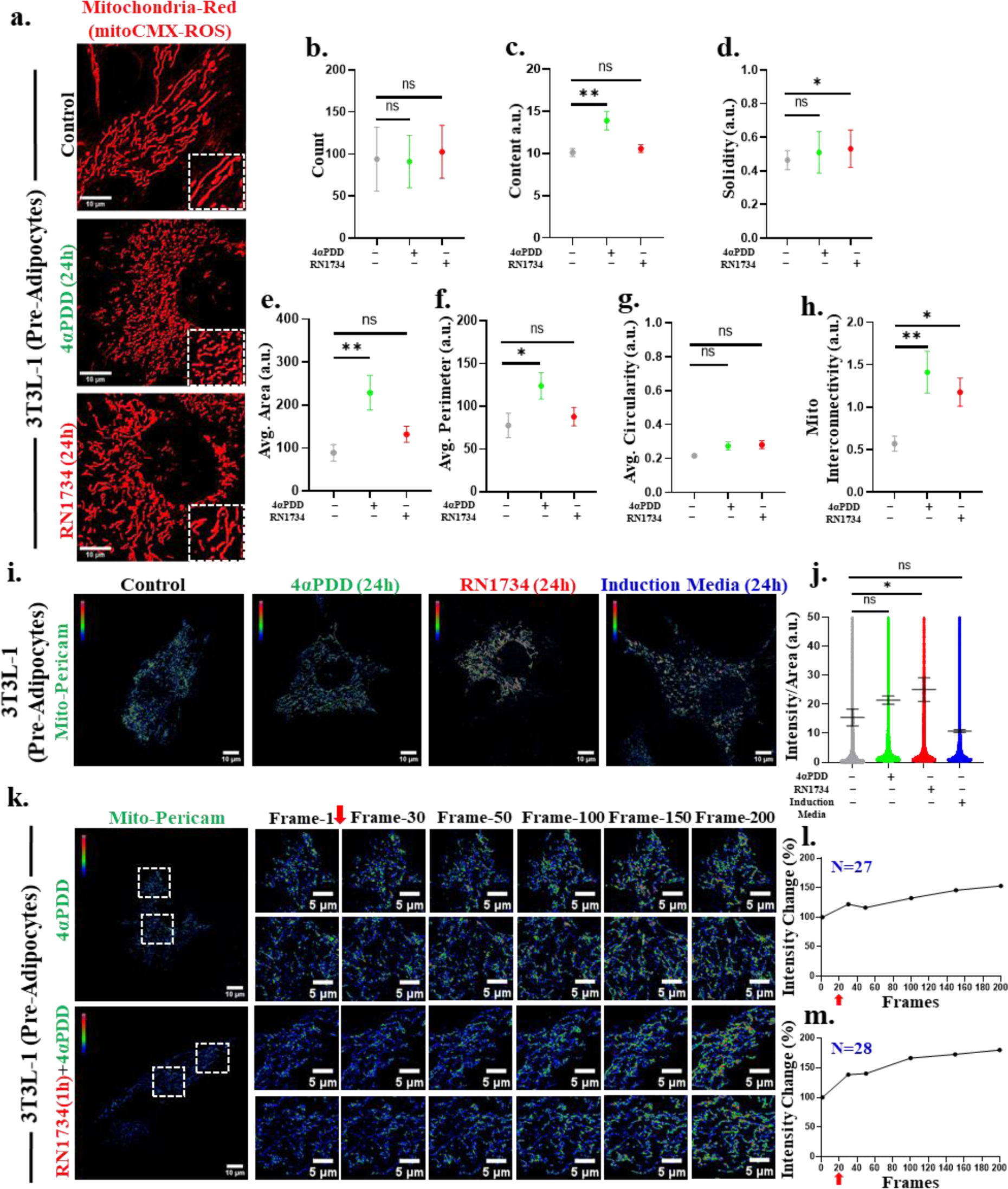
TRPV4 regulates mitochondrial structure and Ca^2+^-dynamics in 3T3L-1 pre-adipocytes. **A.** 3T3L-1 pre-adipocytes were treated with 4αPDD (5μM) and RN1734 (10μM) for 24 hours and stained with MitoTrackerRed-CMXRos. Confocal images were acquired and analysed using Mito morphology plugin in ImageJ. **B-H.** Shown are the quantitative analysis of various mitochondrial parameters. TRPV4 modulation does not affect the mitochondrial count and circularity. TRPV4 activation increases the mitochondrial content, area, perimeter and inter-connectivity. RN1734 leads to increase in mitochondrial solidity and inter-connectivity. n>30 cells (>2000 individual mitochondria) in each conditions were measured. One-way ANOVA; ns =non-significant; * = p<0.05; ** = p<0.01. Scale bar 10μm. **I-J.** Long-term inhibition of TRPV4 leads to more mitochondrial Ca^2+^. Cells were transfected with MitoPericam (mitochondria-specific Ca^2+^-indicator) and subsequently treated with 4αPDD (5μM), RN1734 (10μM) and induction media for 24 hours. n>15 cells (>10000 mitochondria particles). Shown are the intensity profile images in rainbow RGB scale. One-way ANOVA, * = p<0.05; ns = non-significant. Scale bar 10μm. **K.** TRPV4 activation causes instantaneous mitochondrial Ca^2+^-influx. Live cell imaging for mitochondrial Ca^2+^ was performed for 200 frames (∼7.5mins) and 4αPDD (5μM) was added at 20th frame of imaging. In RN1734 + 4αPDD condition, cells were pre-treated with RN1734 for 1 hour. **L-M**. Addition of 4αPDD leads to instantaneous Ca^2+^-influx in mitochondria in both the conditions, however, the extent of Ca^2+^ intake is more in RN1734+4αPDD condition. White box represents the digital zoom. More than 15000 individual mitochondrial particles were quantified in each conditions. Scale bar 10μm; and 5μm for zoom images.

### Long-term inhibition of TRPV4 raises the mitochondrial Ca^2+^

As long-term inhibition of TRPV4 increases the intra-cellular Ca^2+^, we investigated the effect of long-term modulation of TRPV4 (for 24 hours) on mitochondrial Ca^2+^ using MitoPericam as a probe (**Fig 5i**). In case of 4αPDD-treatment, a non-significant change in mitochondrial Ca^2+^-level is seen. But in case of RN1734-treatment, the MitoPericam intensity rises up significantly which depicts the rise in the mitochondrial Ca^2+^ of these cells (**Fig 5j**). Only induction media treatment for 24 hours showed a non-significant decrease in the mitochondrial Ca^2+^ (**Fig 5j**). Results suggest that the long-term inhibition of TRPV4 elevates both mitochondrial and cellular Ca^2+^-levels.

### TRPV4 activation increases mitochondrial Ca^2+^ and decreases mitochondrial particle numbers

We tested the immediate effect of modulation of TRPV4 on mitochondrial Ca^2+^. We performed live cell imaging using the MitoPericam construct for 200 frames (∼7.5mins) and added 4αPDD at the 20^th^ frame (**Fig 5k**). We used a particle-based quantification approach to quantify the mitochondrial particle numbers and their corresponding intensity of different time points (Frame 1, 30, 50, 100, 150 and 200) as described before (Acharya et al 2022b). In 3T3L-1 pre-adipocytes, upon addition of 4αPDD, sudden increase in the mitochondrial Ca^2+^-level is observed (**Fig 5l**). Additionally, we incubated the 3T3L-1 cells with RN1734 for 1 hour and then added 4αPDD. Addition of 4αPDD in RN1734 pre-treated condition (equivalent to TRPV4-inhibited state) leads to more mitochondrial Ca^2+^-influx as compared to normal condition (**Fig 5m**). We measured the extent of Ca^2+^-influx (Frame 200^th^ – Frame 01^st^) and fold-change of the average value in MitoPericam intensity in these conditions. The change in Ca^2+^-level and also the “fold change” is more in cells treated with RN1734 as compared to only 4αPDD (**Fig 6a&b**). Overall, our data suggests that the activation of TRPV4 by 4αPDD causes a sudden increase in mitochondrial Ca^2+^ in 3T3L-1 pre-adipocytes, but in a context dependent manner.

**Figure 6:**
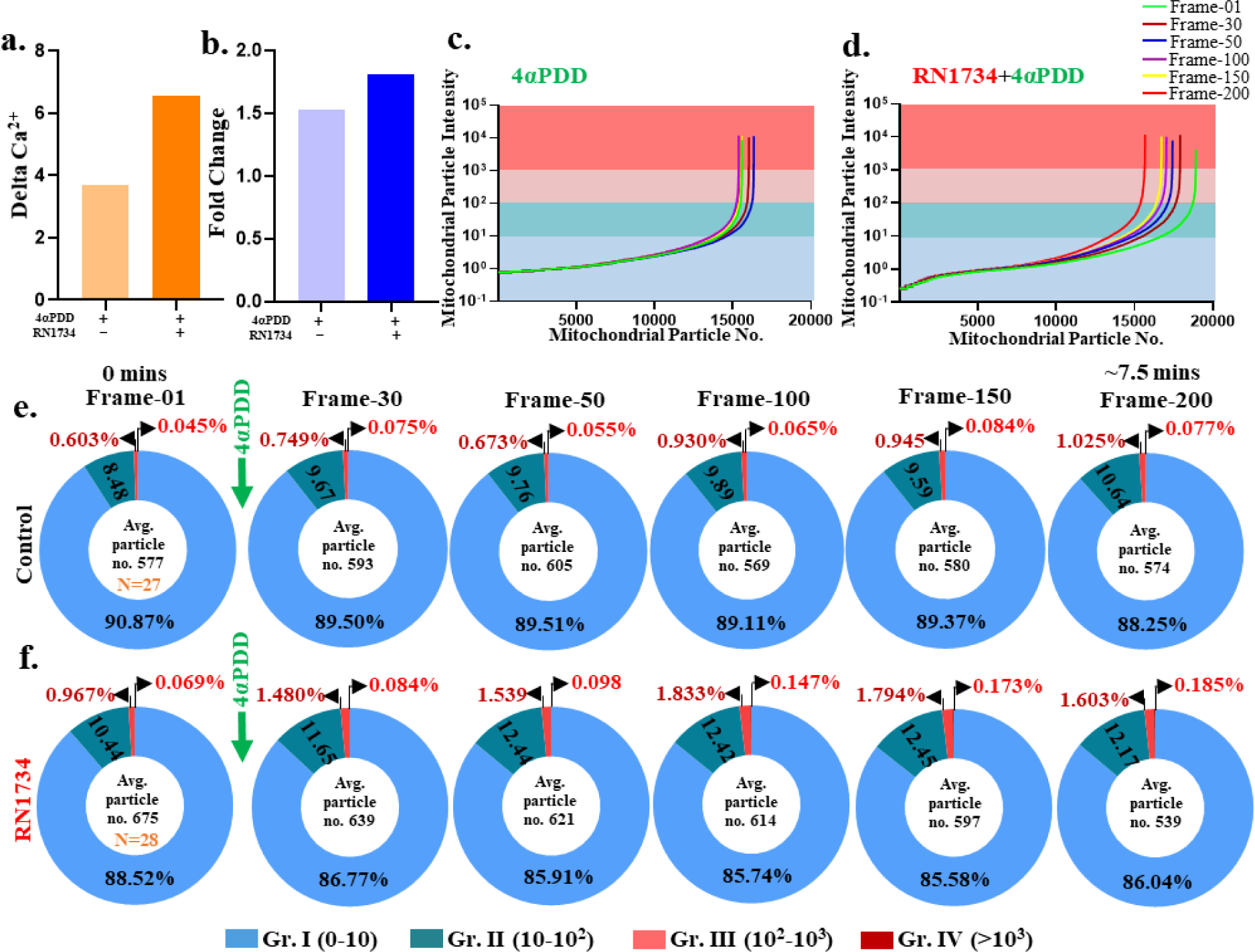
TRPV4 causes instant mitochondrial Ca^2+^-influx and changes in mitochondrial particle numbers. 3T3L-1 pre-adipocytes were transfected with Mito-Pericam and mitochondrial Ca^2+^ imaging was done for 200 frames (∼7.5mins). 4αPDD was added at 20^th^ frame. **A.** RN1734 pre-treatment raises the extent of mitochondrial Ca^2+^-influx instantly as indicated by change in MitoPericam intensity. **B.** Fold change in mitochondrial Ca^2+^ is also more in RN1734 pre-treated cells. **C-D.** TRPV4 activation increases mitochondrial Ca^2+^ instantly and reduces mitochondrial numbers. Upon addition of 4αPDD; the mitochondrial particle reduces. The extent of reduction in number is more in RN1734 pre-treated cells. We classified the mitochondrial particles on the basis of their mitochondrial Ca^2+^ in four groups: Gr I as very low Ca^2+^ (0-10 a.u.; light blue); Gr II as low Ca^2+^ (10-100 a.u.; dark blue); Gr III as high Ca^2+^ (100-1000 a.u.; dark red); and Gr IV as very high Ca^2+^ (>1000 a.u.; bright red). In case of RN1734 pre-treatment, there are more mitochondrial particles that have very less Ca^2+^. **E-F.** Upon activation of TRPV4, the mitochondrial particles with low, high and very Ca^2+^ increases and that of very low Ca^2+^ decreases. Also, the mitochondrial particle number increases in case of RN1734 pre-treatment than control condition.

We next plotted the mitochondrial particle number in an ascending order of their intensities of different time points. Mitochondrial particle number changes upon activation of TRPV4. In case of 4αPDD-treatment, the particle number first increases (during frame 30^th^ & 50^th^) and then decreases (in frame 100^th^ to 200^th^) (**Fig 6c**). Cells treated with RN1734 show mitochondrial particles with lesser mitochondrial Ca^2+^ than control cells (**Fig 6d**). Also, addition of 4αPDD decreases the mitochondrial particle number gradually till the end of the experiment (200^th^ Frame). We also classified the mitochondrial particles in four groups, based on their individual Ca^2+^-intensities as: Gr I: mitochondrial particles with very low Ca^2+^ (0-10 a.u.; light blue); Gr II: with low Ca^2+^ (10-100 a.u.; dark blue); Gr III: with high Ca^2+^ (100-1000 a.u.; dark red); and Gr IV: with very high Ca^2+^ (>1000 a.u.; bright red). In these conditions, the average number of mitochondria decreases upon 4αPDD addition and the percentage of mitochondrial particles with low, high and very high Ca^2+^ increases (**Fig 6e&f**). Also, the mitochondria with very low Ca^2+^ decreases in both these conditions upon 4αPDD addition (**Fig 6e&f**). In summary, the results suggest that TRPV4 activation suddenly increases the mitochondrial Ca^2+^-level and also changes mitochondrial number.

### TRPV4 antagonist RN1734 elevates the mitochondrial superoxide levels during adipogenesis

We explored if and how modulation of TRPV4 affect mitochondrial functions, especially during the adipogenesis. We evaluated the mitochondrial superoxide production using MitoSOX-Red dye (**Fig 7a**). In case of 3T3L-1 pre-adipocytes, 4αPDD results no significant changes, but RN1734 addition leads to a sharp rise in the mitochondrial superoxides (∼2.36 fold) (**Fig 7b**). Similarly, for induced adipocytes also, 4αPDD causes a non-significant increase whereas RN1734 increased the superoxide levels to a great extent (**Fig 7c**). Notably, the basal mitochondrial superoxide level is elevated (∼2.84 fold) in induced adipocytes than pre-adipocytes. Further, in mature adipocytes, the basal mitochondrial superoxide level rises even more (∼4.94 fold) than induced adipocytes. Here too, inhibition of TRPV4 by RN1734 results in more (∼2.19 fold) mitochondrial superoxide (**Fig 7d**). Altogether, the results show that inhibition of TRPV4 by RN1734 during the process of adipogenesis leads to more mitochondrial superoxides. The basal mitochondrial superoxide level increases with differentiation of adipocytes.

**Figure 7:**
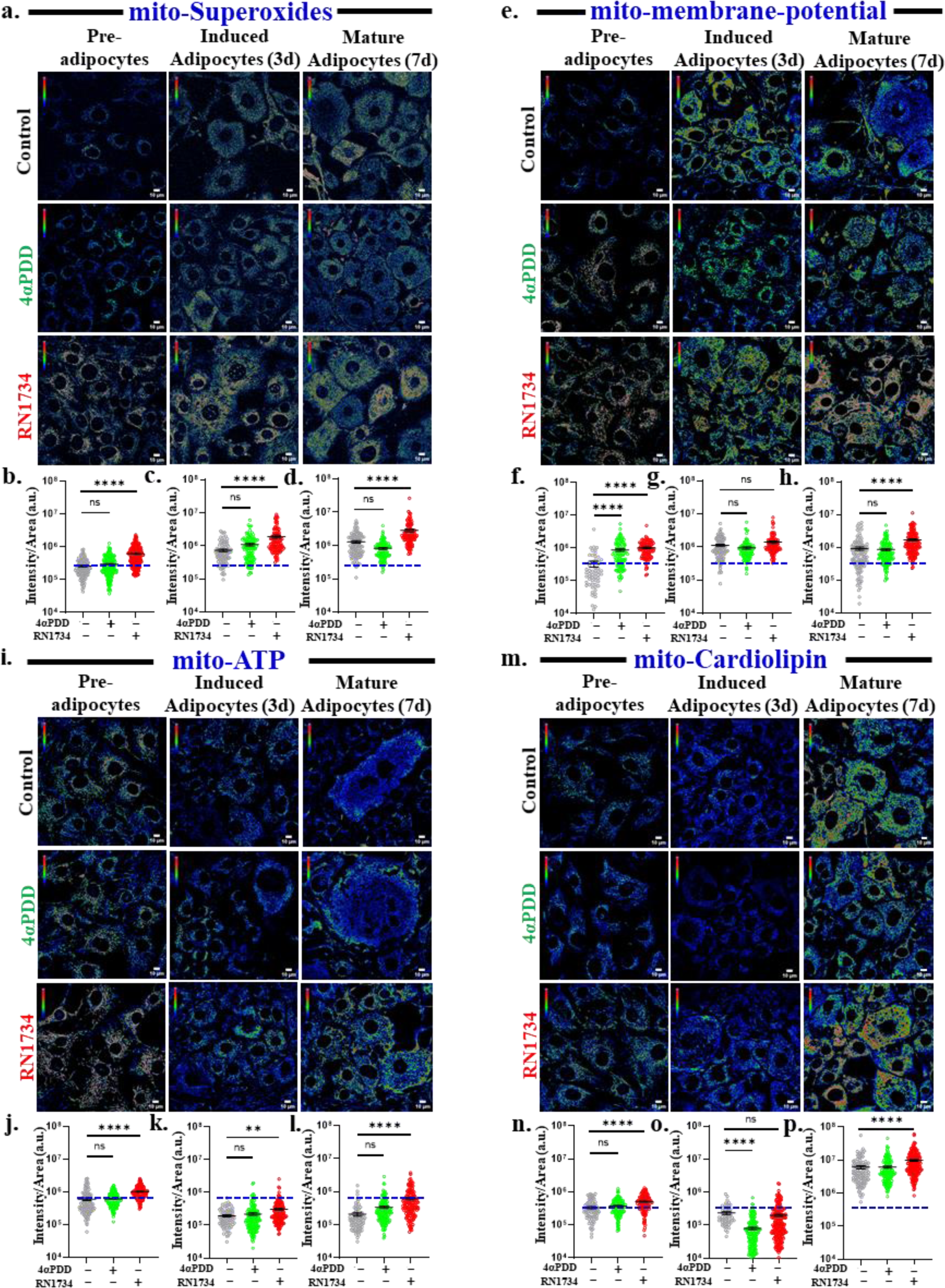
TRPV4 modulation affects mitochondrial metabolism and function during adipogenesis. 3T3L-1 cells were differentiated along with TRPV4 agonist 4αPDD (5μM) and antagonist RN1734 (10μM). For pre-adipocytes, only treatment of modulators was given for 24 hours. For induced adipocytes, images were captured at day 3 of differentiation. For mature adipocytes, differentiation was done for 7 days and subsequently live cell confocal images were taken. Shown are the intensity profile images in rainbow RGB scale of each condition. **A**. TRPV4 inhibition enhances mitochondrial superoxides. MitoSOX Red (2.5μM) was used for 30 mins to label mitochondrial superoxides. **B-D**. Basal level of mitochondrial superoxides increases with adipogenesis. Further, RN1734 addition leads to more superoxides in adipocytes in each condition. **E**. TRPV4 regulates mitochondrial membrane potential. TMRM dye (50μM) was used for 30 mins for detection of mitochondrial membrane potential. **F.** In pre-adipocytes, both activation and inhibition of TRPV4 boosts up the mitochondrial membrane potential. **G.** In induced adipocytes, the basal mito membrane potential enhances. **H.** RN1734-treated mature adipocytes show further increment of mitochondrial membrane potential. **I.** TRPV4 antagonist RN1734 elevates the mitochondrial ATP production. ATP-Red (2.5μM) dye was used for 15 mins to stain mitochondrial ATP. **J.** 3T3L-1 pre-adipocytes enhances ATP production upon RN1734 treatment. **K.** The basal ATP levels reduces in induced adipocytes and then remains almost unchanged in mature adipocytes. **K-L.** In both induced and mature adipocytes, RN1734 causes more ATP levels. **M.** TRPV4 affects mitochondrial cardiolipin levels during adipogenesis. NAO dye (0.5 μM) was used for 15 mins to stain mitochondrial cardiolipin. **N.** Inhibition of TRPV4 by RN1734 causes increment in cardiolipin in pre-adipocytes. **O.** Basal cardiolipin reduces in induced adipocytes. 4αPDD-treated induced adipocytes shows more reduction in cardiolipin. **P.** The basal level of cardiolipin goes up in mature adipocytes. RN1734-treated mature adipocytes shows further increase in the cardiolipin level. n>100 cells in each condition. One-way ANOVA, ** = p<0.01; **** = p<0.0001; ns: non-significant. Scale bar 10μm.

### TRPV4 modulation affects mitochondrial membrane potential during adipogenesis

We next evaluated the dependence of mitochondrial membrane potential on TRPV4 during adipogenesis. We used TMRM dye (which is a mitochondria specific dye that intensifies signal with increment in mitochondrial membrane potential) (**Fig 7e**). In case of 3T3L-1 pre-adipocytes, treating the cells with either 4αPDD or RN173 for 24 hours leads to a significant increase in the mitochondrial membrane potential (**Fig 7f**). In case of induced adipocytes, the basal mitochondrial membrane potential increases than the pre-adipocytes (∼3.4 fold). But modulation of TRPV4 does not affect the mitochondrial membrane potential in induced adipocytes, although the intensity increases non-significantly in RN1734-treated condition (**Fig 7g**). In mature adipocytes, the basal mitochondrial membrane potential slightly decreases (∼0.8 fold) than the induced adipocytes but is greater than pre-adipocytes (∼2.75 fold). TRPV4 activation by 4αPDD during adipogenesis shows no change in mitochondrial membrane potential (**Fig 7h**). RN1734 application during adipogenesis results in more mitochondrial membrane potential of mature adipocytes (**Fig 7h**). Taken together, our data suggests that TRPV4 modulation during adipogenesis affects mitochondrial membrane potential. The basal mitochondrial membrane potential increases in committed step, i.e. in induced adipocytes which then lowers in mature adipocytes.

### Inhibition of TRPV4 during adipogenesis raises the mitochondrial ATP production

We employed ATP Red, a mitochondria-specific dye whose intensity increases with increase in mitochondrial ATP levels (**Fig 7i**). In pre-adipocytes, 4αPDD addition shows no changes in the mitochondrial ATP whereas, RN1734 addition boosts up the mitochondria ATP levels greatly (**Fig 7j**). In case of induced adipocytes, the basal mitochondrial ATP level decreases significantly as compared to pre-adipocytes (**∼**0.32 fold). Further, activation of TRPV4 has no effect on the ATP level whereas, inhibition raises the intensity which implies the rise in ATP levels of mitochondria (**Fig 7k**). Lastly, in mature adipocytes, the basal mitochondrial ATP is similar to that of induced adipocytes (∼0.34 fold lesser than pre-adipocytes). As seen previously, here too activation of TRPV4 doesn’t affect significantly, whereas inhibition of TRPV4 during adipogenesis leads to a sharp rise in the mitochondrial ATP levels (**Fig 7l**). These results suggest that inhibiting TRPV4 during adipogenesis causes increment in the mitochondrial ATP levels and the basal ATP production decreases during adipogenesis.

### TRPV4 antagonist raises the mitochondrial cardiolipin during adipogenesis

We used NAO dye which increases its fluorescence intensity with increase in mitochondrial cardiolipin amount (**Fig 7m**). In pre-adipocytes, 4αPDD causes non-significant changes, whereas TRPV4 inhibition increases the mitochondrial cardiolipin significantly (**Fig 7n**). In induced adipocytes, the basal cardiolipin level reduces as compared to pre-adipocytes (∼0.7 fold). TRPV4 activation further reduces the cardiolipin level significantly, although RN1734 treatment shows no change in the cardiolipin levels in induced adipocytes (**Fig 7o**). Next in mature adipocytes, the basal cardiolipin level increases (∼18.37 fold) as compared to pre-adipocytes. However, TRPV4 activation does not affect the cardiolipin levels of mature adipocytes, but inhibition resulted in increased NAO intensity (**Fig 7p**). Altogether our findings suggest that TRPV4 inhibition during the process of adipogenesis increases the mitochondrial cardiolipin level.

### TRPV4 inhibition during adipogenesis decreases the mitochondrial temperature

We tested if TRPV4 modulation affects the mitochondrial temperature during adipogenesis. We used the MTY dye which increases its fluorescence intensity with decrease in the mitochondrial temperature (**Fig 8a**). In pre-adipocytes, 4αPDD treatment shows no significant change, but RN1734 results in increment in the MTY intensity suggesting the decrease in the mitochondrial temperature (**Fig 8b**). In induced adipocytes, the MTY intensity decreases (∼0.2 fold) which depicts the rise in mitochondrial temperature in induced adipocytes. 4αPDD causes more reduction in MTY intensity (i.e. more increment in mitochondrial temperature) (**Fig 8c**). In addition, RN1734-treatment shows elevation in MTY intensity (lesser mitochondrial temperature) (**Fig 8c**). However, the mitochondrial temperature reaches its normal level/s in mature adipocytes where the MTY intensity is ∼0.84 fold lesser than pre-adipocytes. Further, inhibiting TRPV4 shows an increase (∼1.57 fold) in MTY intensity (i.e. decrease in mitochondrial temperature) whereas, activation shows no effect in mature adipocytes (**Fig 8d**). Overall, our data suggests that in general, inhibition of TRPV4 in adipocytes leads to decrease in mitochondrial temperature during adipogenesis.

**Figure 8:**
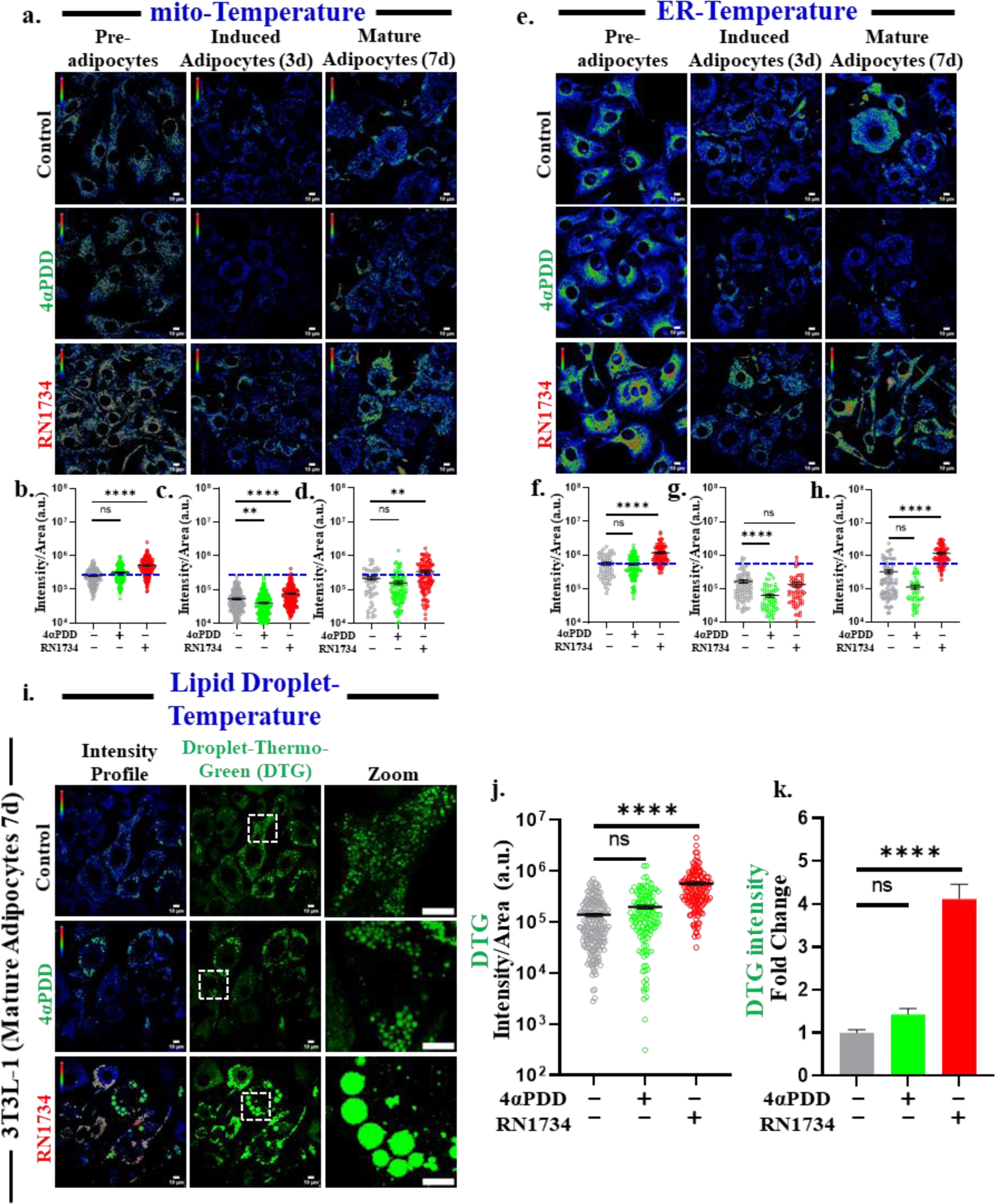
TRPV4 regulates mitochondrial, ER and lipid droplet temperature during adipogenesis. 3T3L-1 cells were differentiated along with TRPV4 agonist 4αPDD (5μM) and antagonist RN1734 (10μM). For pre-adipocytes, only treatment of modulators was given for 24 hours. For induced adipocytes, images were acquired at day 3 of differentiation media before addition of maintenance media. For mature adipocytes, differentiation was done for 7 days and then live cell confocal images were acquired. Shown are the intensity profile images in rainbow scale. Image J was used for quantification. **A.** TRPV4 modulation changes mitochondrial temperature during adipogenesis. MTY dye (0.5μM) was used to assess mitochondrial temperature. Increase in MTY intensity indicates the decrease in mitochondrial temperature. **B.** RN1734-treatment reduces the mitochondrial temperature in pre-adipocytes. **C.** The basal mitochondrial temperature increases in induced adipocytes that further goes up in 4αPDD (decrease in intensity) treated and reduces in RN1734 (increase in intensity) treated conditions. **D.** Mitochondrial temperature become comparable in mature adipocytes and RN1734-treated mature adipocytes show further decrease in the mitochondrial temperature. **E.** TRPV4 regulates ER temperature during adipogenesis. **F.** RN1734 addition reduces the ER temperature in pre-adipocytes. **G.** ER temperature rises during adipogenesis process. 4αPDD increases the ER temperature further in induced adipocytes. **H.** The ER temperature restores in mature adipocytes (slight more than pre-adipocytes) and RN1734 treatment cools the ER during adipogenesis. **I.** TRPV4 inhibition decreases the lipid droplet temperature. DTG (0.5μM) was used to assess the lipid droplet temperature. **J-K.** RN1734-treated cells have less lipid droplet temperature (∼4-fold less). n>100 cells in each condition. One-way ANOVA, ** = p<0.01; **** = p<0.0001; ns: non-significant. White box represents the digital zoom. Scale bar 10μm.

### TRPV4 regulates the ER-temperature during adipogenesis

We used ETY dye which increases its intensity with decrease in ER-temperature (**Fig 8e**). In pre-adipocytes, RN1734 increases the ETY intensity (i.e. decrease in the ER temperature) (**Fig 8f**). In induced adipocytes, the basal level of ETY intensity decreases (∼0.29 fold) indicating the increase in ER temperature. 4αPDD-treated cells show further and significant decrease in the ETY intensity suggesting the presence of relatively hotter ER, whereas RN1734 produces no significant change in the ETY intensity in induced adipocytes (**Fig 8g**). In mature adipocytes, the basal ETY intensity increases than induced adipocytes but remains ∼0.59-fold lesser than pre-adipocytes. Here also, RN1734 makes the ER relatively cold (i.e. more ETY fluorescence intensity) (**Fig 8h**). All in all, TRPV4 affects ER-temperature and ER become relatively hotter during adipogenesis process.

### TRPV4 inhibition during adipogenesis decreases temperature of the lipid droplets

We tested if TRPV4 modulation is affecting the temperature of lipid droplets that are formed during adipogenesis. We employed the Droplet Thermo Green (DTG) probe to label the lipid droplets in mature adipocytes (**Fig 8i**). An increase in the intensity corresponds to decrease in the temperature of the lipid droplets. In case of TRPV4-agonist treated cells, these is a non-significant change in the intensity of the DTG suggesting no major alterations in the temperature of the lipid droplets (**Fig 8j**). However, in TRPV4-antagonist treated cells, the DTG intensity increases significantly (∼4-fold) depicting an overall reduction in the temperature of the lipid droplets in this condition (**Fig 8j&k**). The results depict that inhibiting TRPV4 during adipogenesis causes a drop in the temperature of the lipid droplets that are being formed.

### TRPV4 enhances adipogenesis in primary murine-derived mesenchymal stem cells (MSC’s)

We repeated the major finding in primary MSC’s that were differentiated with or without the modulation of TRPV4. Live cell imaging of BODIPY493/503 was performed to evaluate the adipogenesis process (**Fig 9a**). TRPV4 inhibition during adipogenic differentiation of MSC’s results in elevated levels of intracellular lipids (**Fig 9b**). The BODIPY493/503 intensity increases by ∼2 fold in RN1734-treated condition (**Fig 9l**). TRPV4 activation shows no significant effect on the amount of intracellular lipids (**Fig 9l**). Furthermore, we quantified numbers and morphological parameters of >1000 individual lipid droplets from each condition. TRPV4 activation shows a non-significant increase whereas inhibition slightly reduces the number of lipid droplets per cell (**Fig 9c**). Notably, TRPV4 inhibition leads to an increase in the area, perimeter, radius and volume of the lipid droplets (**Fig 9d-g**). TRPV4 activation during adipogenesis shows reduction in these parameters of the lipid droplets (**Fig 9d-g**). The data suggests that TRPV4 inhibition during adipogenesis upregulates the lipid accumulation in cells and leads to production of bigger lipid droplets.

**Figure 9:**
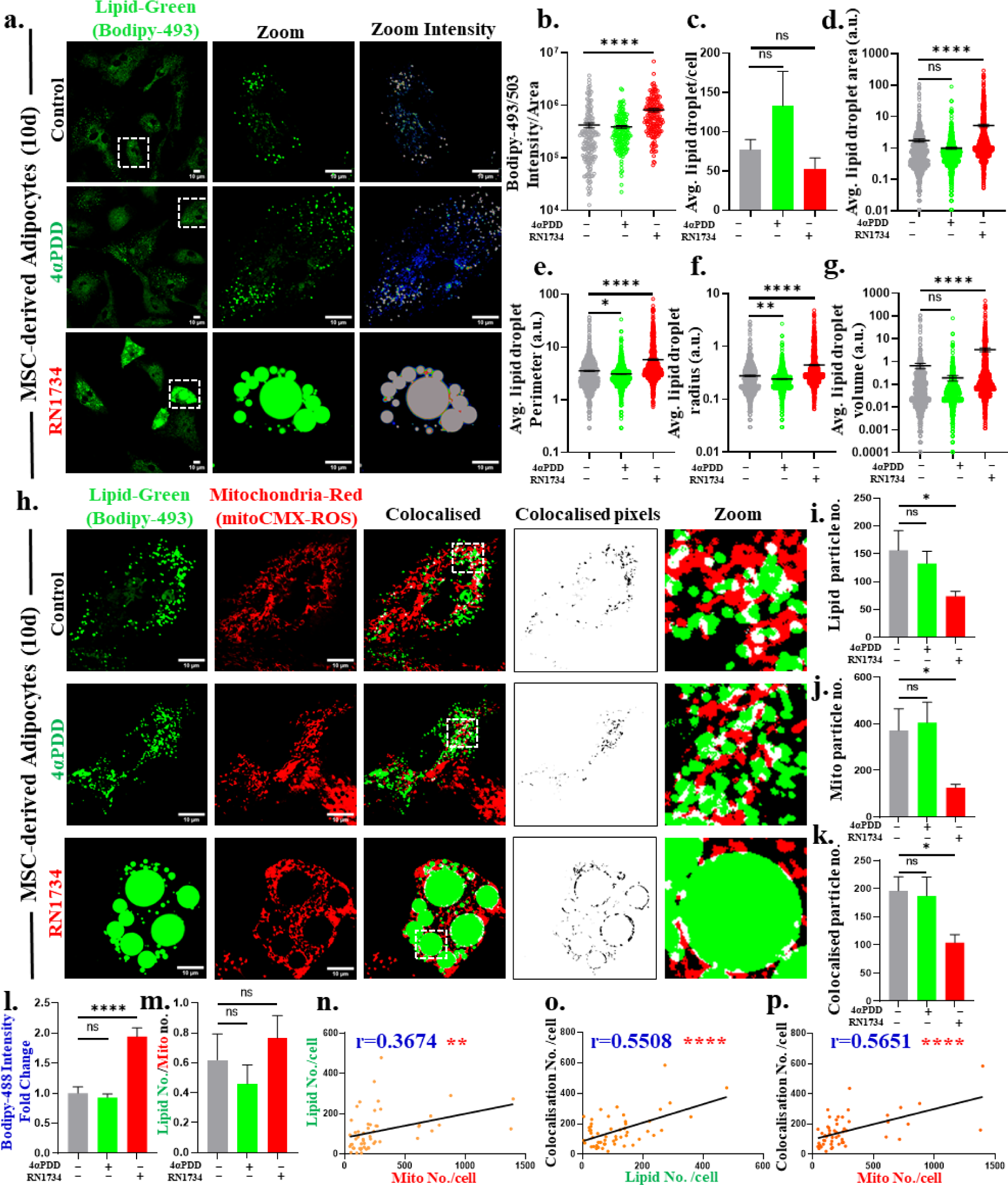
TRPV4 inhibition enhances lipid accumulation in primary murine mesenchymal stem cell-derived adipocytes and decreases the lipid-mitochondria contact site. Primary murine MSC’s were differentiated into adipocytes in presence of either 4αPDD (5μM) or RN1734 (10μM). **A.** Live cell confocal images of adipocytes stained for lipid droplets with BODIPY 493/503 are shown. **B.** TRPV4 inhibition (by RN1734) enhances the lipid accumulation in primary MSC-derived adipocytes. **L.** ∼2-fold increase in intensity of BODIPY493/503 is seen in RN1734-treated adipocytes. **C.** No significant changes were observed in number of lipid droplets/cell in case of TRPV4 activation or inhibition. **D-G.** TRPV4 inhibition during adipogenesis make the lipid droplets bigger as indicated by the increase in area, perimeter, radius and volume of lipid droplets in case of RN1734 treatment. **D-E.** 4αPDD reduces the radius and perimeter of the lipid droplets. n>150 cells in each condition. One-way ANOVA * = p<0.05; ** = p<0.01; **** = p<0.0001; ns: non-significant. Scale bar 10μm. **H.** Shown are confocal images of lipid (green) and mitochondria (red) of primary MSC-derived adipocytes. **I-K.** Analysis of ∼6000 lipid particles and ∼15000 mitochondrial particles in all conditions was carried out. Inhibiting TRPV4 during adipogenesis reduces the lipid numbers, mitochondria numbers, and the lipid-mitochondria contact points. **M.** TRPV4 modulation changes the ratio of lipid/mitochondria number in primary adipocytes. **N-P.** The correlation analysis suggests that the lipid number and mitochondria number are mostly independent on each other. **O-P.** Lipid-mitochondria contract points are almost equally dependent on lipid as well as mitochondrial numbers (similar correlation values). One-way ANOVA, * = p<0.05. Scale bar 10μm.

We plotted the individual lipid droplets in an ascending order of their area. A bi-phasic curve for each of control, 4αPDD and RN1734 condition was observed (**Fig S3b**). Notably, the lowest size of the lipid droplet in RN1734 –treated condition is also bigger than the lowest droplet observed in control or 4αPDD-treated conditions. We used a similar classification for the lipid droplets as defined by the ORO-labelling (**Fig S3d**). Firstly, in RN1734-treated condition, the percentage of small-sized lipid droplets reduces significantly from control condition (68.86% to 45.48%). Also, the small-sized lipid droplets increased in 4αPDD-treated condition (to 78.09%) (**Fig S3d**). The medium-sized lipid droplets increased in RN1734 treated condition (44.78%; dark blue) as compared to control (29.55%) and decreased in 4αPDD-treated conditions (21.17%). Most importantly, the percentage of lipid droplets with large-size (3.5%; light red) and very-large size (6.23%; dark red) is significantly increased in RN1734-treated condition than control where large (0.76%) and very large (0.83%) lipid droplets remained at very low abundance. In 4αPDD-treated condition, the large (0.43%) and very large (0.31%) lipid droplets reduced further than control. In summary, our results and analysis suggest that TRPV4 modulation during adipogenesis changes the pattern of lipid distribution in cells (based on their size) and TRPV4 inhibition causes formation of bigger lipid droplets.

### TRPV4 regulates lipid-mitochondria contact points in primary mesenchymal-derived adipocytes

Since TRPV4 modulation affects the size of the lipid droplets and mitochondria, we tested if TRPV4 modulation affects the lipid-mitochondrial contact points in adipocytes (**Fig 9h**). Primary MSC’s were differentiated with or without the TRPV4 modulators. Total lipid, mitochondria and colocalised particle numbers were quantified. Data suggests that TRPV4 inhibition decreases the total lipid numbers, mitochondrial numbers and the lipid-mito contact points (**Fig 9i-k**). We also observed that in TRPV4-inhibited cells, the lipid-mitochondrial contact points are mostly visible around the lipid droplets (often around the bigger lipid droplets), whereas, in control and TRPV4-activator treated cells the contact points are scattered throughout the cell. We plotted the ratio of lipid/mito numbers of the above conditions. The lipid/mito ratio reduces in 4αPDD-, but increases in RN1734-treated condition (**Fig 9m**). We also analysed the dependency of lipid-mito contact points on lipid and mitochondria by estimating the correlation. There is no strong correlation in lipid and mitochondria numbers (r = 0.3674) (**Fig 9n**). The correlation of lipid-mito contact number with both lipid number (r = 0.5508) and mitochondria number (r = 0.5651) were almost similar and remain at a modest value (**Fig 9o&p**). Overall, our data indicates that the lipid-mito contact point changes via modulation of TRPV4 and TRPV4 inhibition during adipogenesis significantly reduces the mitochondrial number. The lipid-mito contact points mostly remain equally dependent on both lipid numbers and mitochondria numbers.

## Discussion

Obesity is a global problem which is currently turning into an unprecedented epidemic [28]. In obese condition, adipose tissue increases which is governed by the process of adipogenesis manifested in adipocytes [29]. Temperature and adipogenesis are often correlated where both core body temperature and environmental temperature contributes to weight gain [30]. TRPV4 is an important thermosensitive ion channel which is activated at warm temperatures (>34°C) [31]. TRPV4 can also be activated by mechanical-force [32]. TRPV4 is reported to affect the adipose oxidative metabolism; thermogenesis and expression of pro-inflammatory genes *in vivo* [33, 34]. However, the involvement of TRPV4 in the process of adipogenesis is still questionable, remain contradictory and largely unexplored at the cellular and subcellular level. In this work, we have evaluated how TRPV4 regulates the adipogenesis process especially in context of mitochondrial metabolism and functions by using isolated cultured adipocytes. This arguably cut-off the influence of factors secreted by other cells and tissues and eliminates the cross-talks of different signalling. Our work suggests that TRPV4 acts as a key regulator of adipogenesis and is relevant for adipocyte biology.

Pharmacological inhibition of endogenous TRPV4 functions leads to higher level (often also bigger sized) of lipid-droplet formation in both 3T3L-1 cells and murine-derived mesenchymal stem cells. Inhibition of TRPV4 also increases the percentage of cells that become positive for Oil Red O (adipocyte turnover). This suggests that endogenous TRPV4 activation prevents differentiation of pre-adipocytes to mature adipocytes and/or excess lipid droplet formation. If such enhanced lipid droplet is due to more lipid formation and/or less lipid degradation, that remains to be explored in future. Enhanced expression of SCD1 also suggests that inhibition of TRPV4 might also changes the composition of the lipid-droplets [35]. Never-the-less, several reasons suggest that endogenous TRPV4 function might also be relevant for lipid degradation along with lipid synthesis. First, total number of detectable lipid droplets remain more-or-less constant in different conditions, suggesting that TRPV4 activation may not increase the number of lipid-droplets *per se*. Second, comparatives of the “lowest size of the lipid droplets” in different conditions suggest that inhibition of TRPV4 results in increased size of lipid droplets. Third, the data suggests reduction in mitochondrial number and lipid-droplet-mitochondrial contact points due to inhibition of TRPV4. All these aspects make the importance of TRPV4-mediated mitochondrial regulation more relevant.

Expression of different TRPV members, such as TRPV1, TRPV2, TRPV3, TRPV4 & TRPV6 in mouse adipocytes (3T3L-1) was reported [36]. Notably, relative expressions of TRPVs change in pre-adipocytes and mature adipocytes [36, 37]. TRPV4 expression alters during the process of adipogenesis. This study indicates that the expression of TRPV4 increases in mature adipocytes (∼2 fold) as compared to pre-adipocytes. We have not noted any decrease in TRPV4 expression in mature adipocytes as reported previously [36]. Our data also suggests that blocking TRPV4 function by pharmacological agents during adipogenesis increases endogenous TRPV4 level in mature adipocyte.

In this work we demonstrate that TRPV4 regulates a number of sub-cellular parameters, such as remodelling of actin cytoskeleton, mitochondrial distribution and functions during the differentiation of pre-adipocytes to mature adipocytes. TRPV4 also seems to regulate contact points between lipid-droplets and mitochondria, suggesting overall and diverse changes in cell that favours formation of lipid droplets (**Fig 10g**). Direct involvement of TRPV4 also opens scopes to apply TRPV4-specific pharmacological agents for possible reduction of adipocyte differentiation and/or lipid accumulation (data not shown).

**Figure 10:**
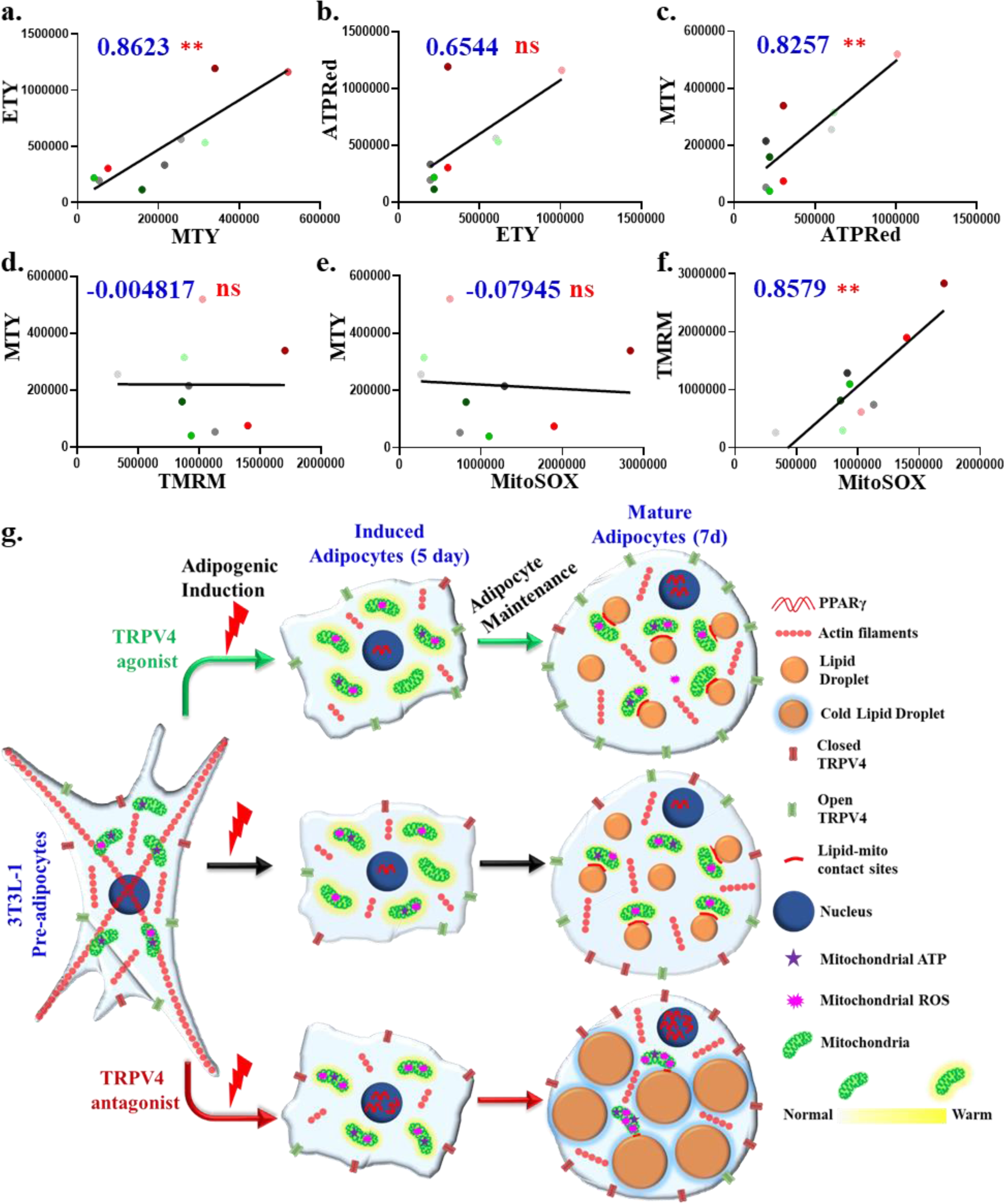
TRPV4-mediated changes in mitochondrial-metabolism are co-related. **A-F.** The average values of the 9 different condition across 3 different stages of adipogenesis (pre, induced and mature adipocytes) were plotted and their co-relation was evaluated. Pearson’s r value was calculated (written in blue). p-value, ** = p <0.01; ns: non-significant. **G**. Schematic model depicting the TRPV4-mediated control of adipogenesis. TRPV4 modulation affects differentiation of pre-adipocytes to mature adipocytes. Inhibition of TRPV4 during adipogenesis leads to formation of bigger lipid droplets, lesser mitochondrial number and lipid-mitochondria contact sites. Expression level of TRPV4 initially decreases in induced-adipocytes and then increases (∼2 fold) in mature adipocytes. Adipogenic induction disrupts the F-actin filaments in pre-adipocytes which is facilitated in TRPV4-antagonist treated cells. Mitochondrial temperature increases with the differentiation of adipocytes which then reduces in mature adipocytes. TRPV4 inhibition reducers temperature of lipid-droplets possibly due to reduction in mitochondrial temperature and reduced physical contact between mitochondria with lipid droplets. Furthermore, mitochondrial ATP level reduces with differentiation while mitochondrial superoxides level increases. In case of TRPV4-antagonist treated adipocytes, the ROS and ATP remain at a higher level with lesser mitochondrial temperature. TRPV4 seems to regulate the adipocyte number, differentiation, metabolism, and turnover.

Cytoskeleton remodelling is critical for the process of adipogenesis [26, 38]. Long and dense-actin fibres induce sufficient mechanical pressure, cell stiffness, shape and bio-mechanics that prevents lipid-droplet formation. Indeed, actin disrupting agents are also reported to induce more adipogenesis, which in turn guides the adipocytic differentiation of cells [26, 39, 40]. Disruption of actin filaments are required for the induction of adipogenesis by stem cells [27]. Our data accords well with the previous finding that shows the levels of F-actin changes during the course of adipogenesis [8]. Here we show that extent of F-actin first decreases and then recovers in mature adipocytes [41]. However, the qualitative differences in terms of the long stress fibre become prominent. TRPV4 is a known interactor of cellular cytoskeleton particularly of tubulin and F-actin [25]. These findings also correlate with the changes in the cellular morphology (size, length, and breadth) of cells during adipogenesis process. Our findings also suggest that the adipogenic induction-mediated changes in F-actin overpower the TRPV4-mediated changes in actin cytoskeleton.

The mechanism for regulation of adipogenesis by Ca^2+^-signalling is complex and is dependent on the mode by which Ca^2+^-level is changed [42]. Our results suggest that although instantaneous TRPV4 activation causes cytosolic Ca^2+^-influx, but, its long-term inhibition also causes rise (∼1.5 folds) in cytosolic Ca^2+^-levels in 3T3L-1 cells. This is most-likely due to the blockage of intra-cellular Ca^2+^-buffering units that harbours TRPV4 (such as mitochondria). We propose that TRPV4-inhibition-mediated changes in cytosolic Ca^2+^ favours adipogenesis of 3T3L-1 cells. It seems that only a subset of 3T3L-1 pre-adipocytes is sensitive to 4αPDD stimulation and two distinct populations of cells resides within; i.e. as “responsive” and “non-responsive” to TRPV4 activation.

Mitochondrial biogenesis, bioenergetics and remodelling are critical for induction of adipogenesis [10]. A decrease in mitochondrial function is shown to effectively supress adipogenic differentiation of cells [12]. Moreover, mitochondrial dysfunction is correlated with obesity and diabetes [43]. Recently we have reported the presence of full-length TRPV4 in mitochondria [16]. TRPV4 activation causes instantaneous mito-Ca^2+^ influx, changes its structure and ER-mitochondrial contact points [16, 18]. Long-term modulation of TRPV4 also alters mitochondrial Ca^2+^, most-likely due to impaired Ca^2+^-buffering, yet differently in different cell types [18]. TRPV4 modulation is shown to affect the mitochondrial ROS, ATP, and membrane potential which are important factors of adipogenesis. Therefore, TRPV4-mediated regulation of mitochondrial metabolism and other properties during adipogenesis seem to be relevant for lipogenesis.

Mitochondria is a major hub for production of ROS which is required for adipocyte differentiation [10]. A moderate level of ROS is shown to enhance adipogenesis, and mitochondria targeted anti-oxidants attenuates adipogenesis [44]. Accordingly, Arabinosylcytosine-mediated increase in adipogenesis is facilitated by rise in mitochondrial ROS production [45]. Notably, we observed that the mitochondrial ROS levels increases during the course of adipogenesis (∼5 fold in mature adipocytes) which rises by another ∼2.2 folds in RN1734-treated mature adipocytes. Thus increased ROS production and adipogenesis due to RN1734 treatment as shown in this study accords well with previous reports.

Both extra- and intra-cellular ATP levels are shown to modulate adipogenesis [15, 46]. Although mitochondria becomes metabolically active during adipogenesis however a reduction in cellular ATP is reported [47]. But, the corresponding changes in mitochondrial ATP is yet to be explored. Our results indicate a decrease in the mitochondrial ATP during adipogenesis (∼0.3 fold less than pre-adipocytes). The relative increment in mitochondrial ATP in RN1734-treated mature adipocytes indicates their high metabolically active state which can facilitate the adipogenesis process [48].

Cardiolipin are mitochondrial membrane phospholipids that govern the normal function of mitochondria [49]. In adipocyte, they are also involved in the stimulation of thermogenesis [14]. Alterations in the cardiolipin levels are prevalent in obese patients in an organ-specific manner [50]. Our study for the first time shows that amount of cardiolipin is regulated during adipogenesis and ∼18-fold increase in cardiolipin occurs in mature adipocytes than pre-adipocytes. This huge increase might be the consequence of rise in mitochondrial metabolism and mass of mature adipocytes [51]. The cardiolipin changes also point towards the “browning of the adipocytes” as lipidomics of WAT shows no presence of cardiolipin [50]. Mitochondrial membrane potential is an important indicator of mitochondrial health and activity [52]. Previous findings (human MSCs) suggest that mitochondria depolarises upon adipocytic differentiation [12, 47]. However, our data differs from these findings and an hyperpolarisation of mitochondrial membrane (∼2.8 fold in mature adipocytes) was observed in 3T3L-1-derived adipocytes. This variability might be due to the different origins and components (Rosiglitazone and Indomethacin) used for the adipocytic induction of cells. Furthermore, RN1734 makes the mitochondria more active which facilitates the increased adipogenesis of these cells.

Mitochondrial temperature is a by-product of the OXPHOS which is intimately linked with the mitochondrial dysfunctions [53]. It has been reported that “Mitochondria are physiologically maintained at close to 50°C”, suggesting their warmer temperature than the surrounding [54]. We proposed that TRPV4 is able to regulate mitochondrial temperature [16]. Using a similar approach, we identified that mitochondria becomes hot with adipogenic induction and restores its temperature in mature adipocytes. Also TRPV4 modulation affects ER-temperature, at least in certain conditions. Our data depicts rise in ER-temperature during adipogenic induction. So it seems that “temperature gradient” between ER and mitochondria is regulated by TRPV4 to some extent (**Fig 10a**). This “temperature gradient” might be also relevant for ATP-flow from mitochondria to ER (**Fig 10b-c**). However, the mitochondrial temperature seems to be not correlated with the mitochondrial membrane potential and superoxides levels (**Fig 10d&e**). But, we observed that mitochondrial membrane potential and superoxides are highly correlated (**Fig 10f**). We also report a change in the lipid droplet temperature by TRPV4 modulation during adipogenesis. TRPV4 inhibition drastically reduces the lipid droplet temperature, possibly making the lipid droplets “more viscous” (due to lower temperature) which will further decrease the ability of lipid droplets to transfer the fatty acids via the lipid-mitochondrial contact sites [55]. In other words, the data suggests that in normal condition, mitochondria-produced heat energy may be helpful to reduce the “viscosity” (other physico-chemical kinetics) of the lipid-droplets for efficient transfer of fatty acids directly to mitochondrial for β-oxidation. This also points towards the “temperature gradient” between lipid droplets and mitochondria which is required for efficient transfer of fatty acids. Our data also suggests that both lipid droplet and mitochondrial temperature are tightly controlled by TRPV4 and a positive co-relation exits between both (data not shown).

Cholesterol is known to induce change in cell stiffness and guide adipocyte differentiation [56]. Both cholesterol depletion as well as enrichment affects the extent of adipocyte differentiation [56]. TRPV4 is reported to interact with cholesterol which regulates the functionality of TRPV4 [57]. Our data matches well with the previous report which suggests enrichment of cholesterol positively correlates with the adipogenic differentiation as seen in case TRPV4-antagonist-treated cells. Notably, TRPV4 function is also regulated by cholesterol and cholesterol reduction often increases the mobility of TRPV4 [58]. There are several conserved and putative cholesterol-binding motifs present on TRPV4 which gets affected in cases of TRPV4-mediated channelopathies [8, 57, 59, 60]. Also, the syntenic locus of TRPV4 suggests that TRPV4 has co-evolved with cholesterol biosynthesis pathway, at least to some extent in vertebrates [8]. Another mechanosensor (Piezo1) was demonstrated to regulate cholesterol biosynthesis pathway [61]. Our findings are consistent with this notion and suggest blocking of TRPV4 (and/or other mechanosensors) may upregulate the cholesterol level of cells. Moreover, the amount of cholesterol determines the membrane fluidity with may in turn affect other direct or indirect effect in adipogenesis [62, 63]. Mechano-sensation and adipogenesis are interconnected [64]. Several reports suggest the involvement of mechano-sensitive (Piezo) channels in regulation of lipogenesis and adipogenesis [65, 66]. Furthermore, mechano-sensation of cells is critical in determining adipocyte size, number, and adipogenic potential [67]. Lipid accumulation also changes the rigidity and stiffness of adipocytes as measured by AFM [68, 69]. Thus, disrupting the mechanosensation of cells (as in case of RN1743) may lead further lipid accumulation. Moreover, lipid-mitochondria contact sites are demonstrated to be both functionally and structurally critical for lipid droplet size regulation [70, 71]. Exercise-mediated reduction in body weight was accompanied by increased lipid-mitochondria contact sites as seen in muscle biopsy samples [72]. Our results suggest that decrease in mitochondrial content can facilitate the increase in size of lipid droplets. This finding does not accords with previous report which suggests increased lipid-mitochondria are associated with increasing lipid droplet size [73]. However, this might possibly be explained by comparing the bioenergetics of mitochondria in RN1734 which remains unexplored.

We conclude that TRPV4 is directly involved in differentiation of pre-adipocytes to mature adipocytes in isolated cultured condition and lipid accumulation in mature adipocytes. TRPV4 seems to affect the status of F-actin, mitochondrial metabolism-related properties and mitochondrial temperature, all of which can affect adipogenesis process (**Fig 10g**). Our data also suggests for possibilities of altered lipid profile in patients having different mutations in TRPV4 gene. These findings not only increase our knowledge of complex regulation of adipogenesis process particularly by TRPV4, but also highlights the possible application of TRPV4 activators in treating obesity.

## Acknowledgement

This work is supported by intramural grants to C.G by Department of Atomic Energy (DAE, NISER). SK1, SK2 and TKA are supported by scholarship provided by DAE India and University Grant Commission (UGC, India). Authors acknowledge all the present and former lab members for their support and critical comments.

## Author contributions and declaration

CG conceptualized the project, provided reagents, facilities and guidance. SK1, SK2 and TKA performed all the experiments and analysed the data. SK1, SK2, TKA and CG critically analysed the data. SK1, SK2 and TKA compiled the figures for the manuscript. All authors contributed in the manuscript preparation. YTC provided the MTY and ETY dye. CG edited the final manuscript and finalized the author list.

## Conflict of interest statement

All authors declare the existence of no conflict with this work. The funding bodies have no role in experiment design, data analysis, manuscript preparation or decision to publish this work.

## “Data availability” statements

All data generated or analysed in this study are included in this manuscript. The data set can be available from the corresponding author upon reasonable request.

## Abbreviation

3T3L-1: Murine pre-adipocyte cell line
ER: Endoplasmic Reticulum
ETY: ER Thermo Yellow
MA: Mature adipocyte
MTY: Mito Thermo Yellow
NAO: Nonyl Acridine Orange
ORO: Oil Red O
ROS: Reactive Oxygen Species
TRPV4: Transient Receptor Potential Vanilloid sub-type 4
TMRM: Tetramethylrhodamine, methyl ester

## Supplementary figures

**Figure S1:**
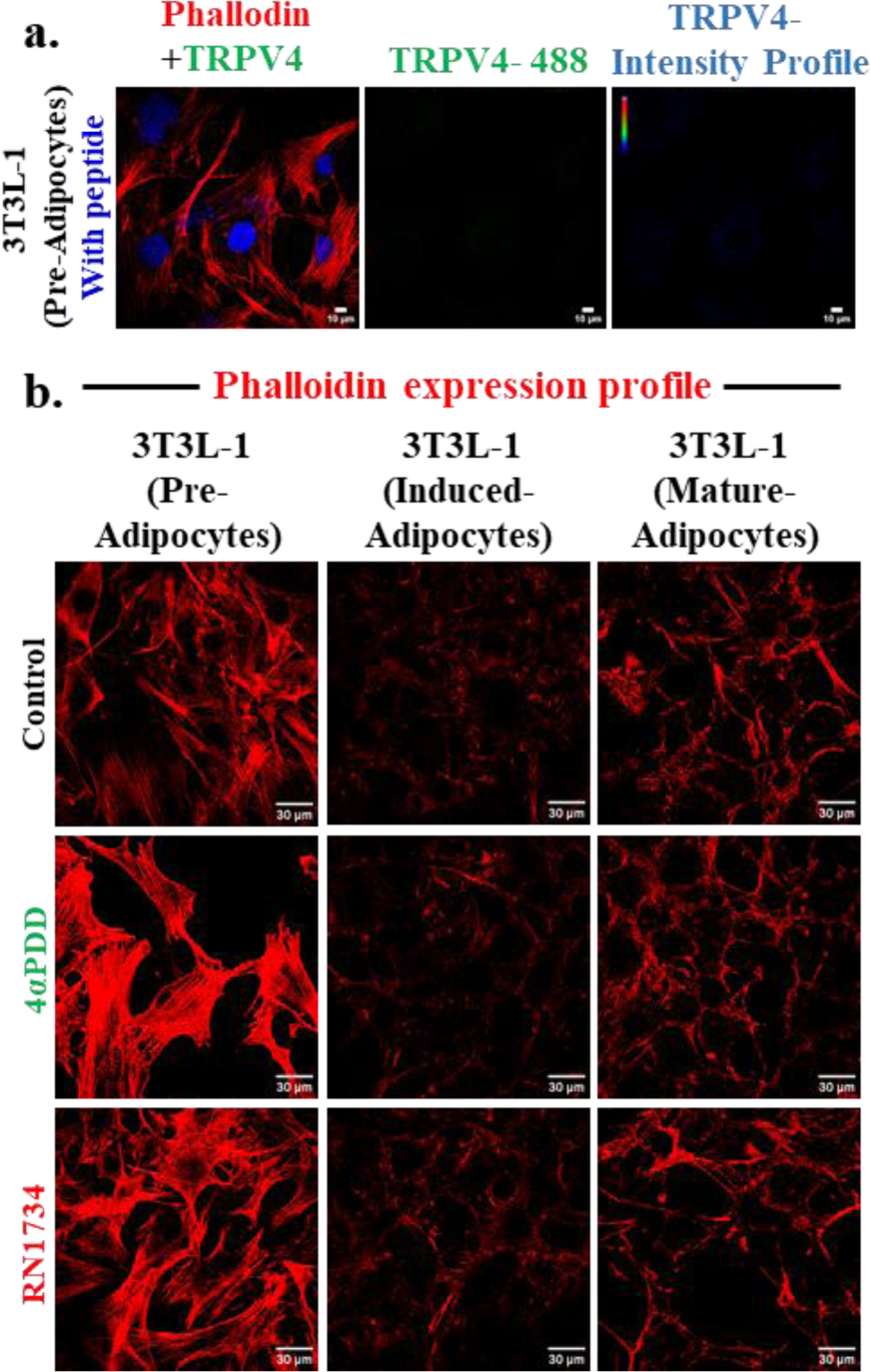
**A.** 3T3L-1 were stained for TRPV4 along with its blocking peptide. Presence of peptide drastically diminishes the TRPV4 specific signal. Scale bar 10μm. **B.** 3T3L-1 cells were differentiated in presence of 4αPDD and RN1734 and stained for Phalloidin to label the F-actins. Shown are the representative images from each condition. Modulation of TRPV4 increases F-actin filaments in pre-adipocytes. In case of induced adipocytes, the F-actin reduces which then increases in mature adipocytes. Scale bar 30μm.

**Figure S2:**
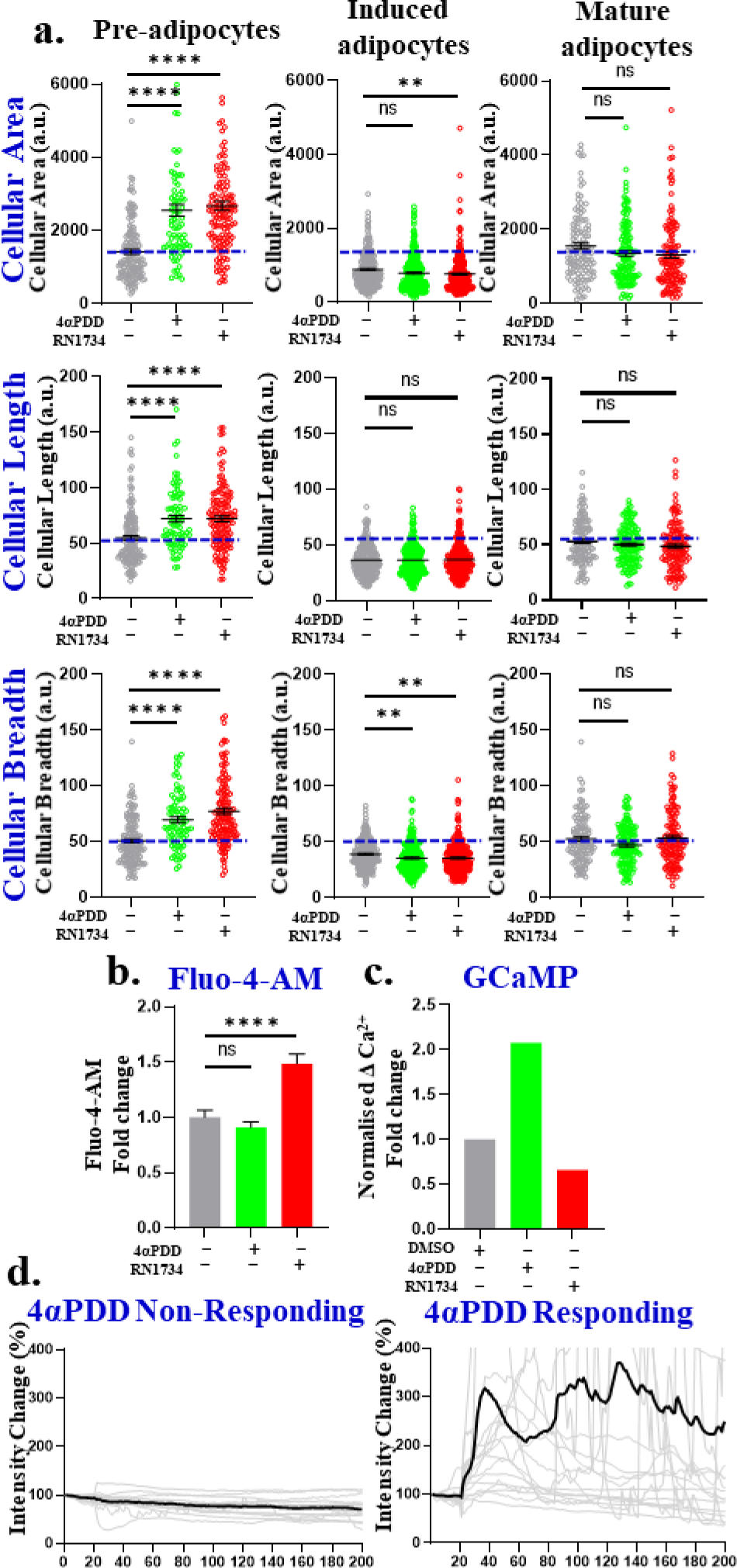
**A. TRPV4 affects cellular morphology during adipogenesis process.** 3T3L-1 pre-adipocytes were differentiated with 4αPDD (5μM) or RN1734 (10μM). For pre-adipocytes, modulation was performed for 24 hours. The cells were stained with Phallodin to measure the morphometric parameters. 4αPDD and RN1734 increases the cellular area, length and breadth in pre-adipocytes. These parameters decrease in induced adipocytes (day 3) (cells become small) and then again increase in mature adipocytes (day 7). In mature adipocytes, TRPV4 modulation does not change the cellular morphological parameters. One-way ANOVA, ** = p<0.01; **** = p<0.0001; ns = non-significant. **B.** TRPV4 inhibition for 24 hours in 3T3L-1 pre-adipocytes increases the cellular Ca^2+^ ∼1.5 fold. **C.** Instant activation of TRPV4 in 3T3L-1 pre-adipocytes raises the cellular Ca^2+^ by ∼2 fold, whereas inhibition reduces it by ∼0.7 fold as seen by GCaMP intensity change. **D.** In the GCaMP transfected pre-adipocyte cells, two distinct cell groups exist, i.e. “responding” and “non-responding” when TRPV4 is activated by 4αPDD. The responsive cells have huge cellular Ca^2+^-influx whereas, non-responding cells are not affected by TRPV4 activation instantly.

**Figure S3:**
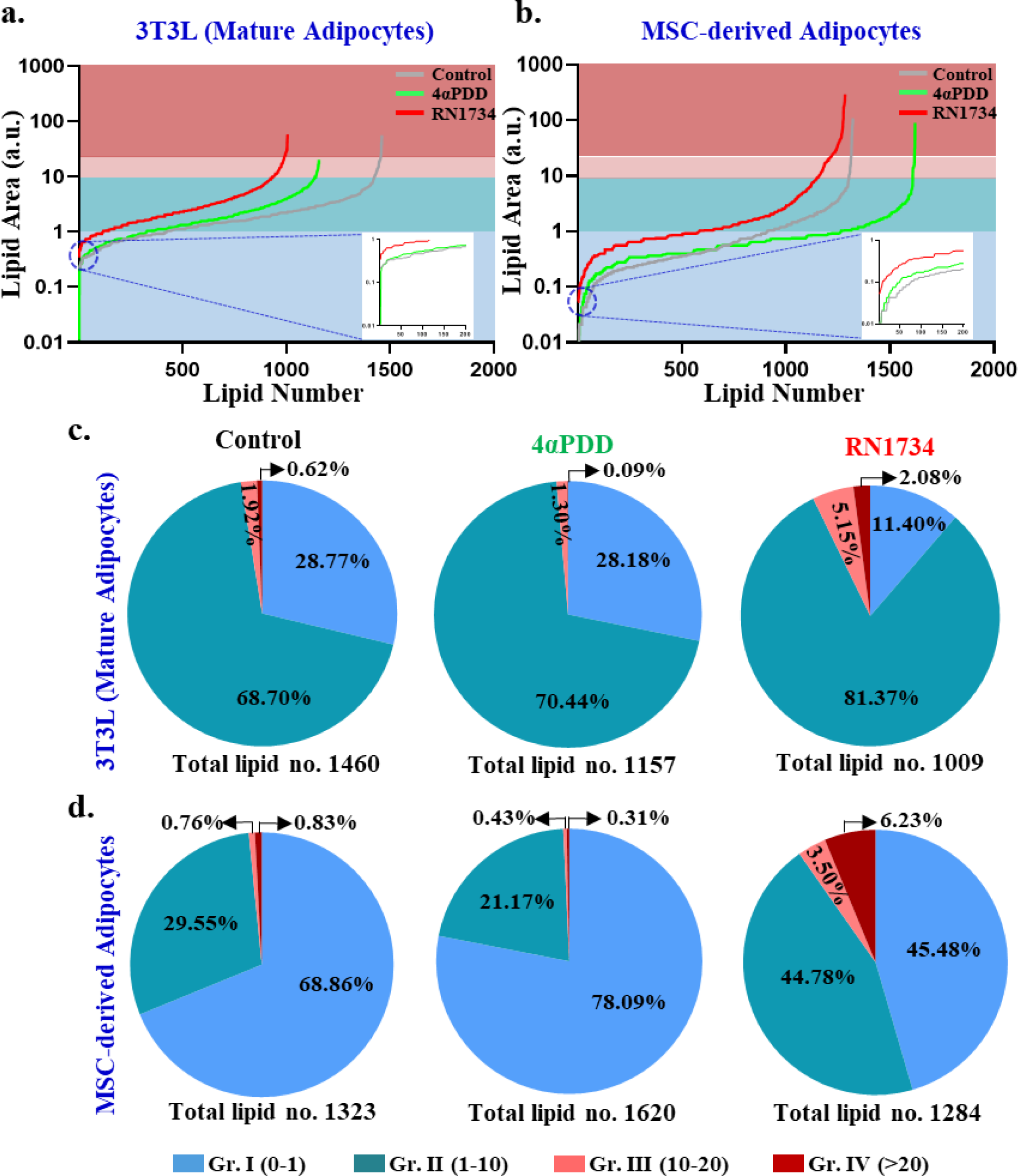
TRPV4 modulation during adipogenesis changes the numbers and sizes of the lipid droplets. **A.** 3T3L-1 cells were differentiated in the presence of 4αPDD or RN1734. Modulation of TRPV4 reduces the lipid droplets number slightly. For RN1734, the line starts from above suggesting high proportion of bigger lipid droplets (inset). **B.** Primary MSC-derived adipocytes (in presence of TRPV4 modulators). Here also, the lipid droplets number changes in 4αPDD (increases) and RN1734 (decreases) condition. Also, the size of the droplet is biggest in RN1734 (red line goes highest). **C-D.** TRPV4 changes lipid droplet percentage on the basis of their sizes. We classified various lipid droplets on the basis of their size (area) as small (0-1 a.u, light blue); medium (1-10 a.u., dark blue); large (10-20 a.u., light red); and very large (>20 a.u, dark red). In both MSC and 3T3L-derived adipocytes, in RN1734 condition, the percentage of medium, large and very large lipid droplet increases as compared to control and that of small lipid droplet decreases. In 4αPDD, the large and very large lipid droplets percentage decreases.

## Notes

### Competing Interest Statement

The authors have declared no competing interest.

## References

[1] J.P. White, M. Cibelli, L. Urban, B. Nilius, J.G. McGeown, I. Nagy, TRPV4: molecular conductor of a diverse orchestra, Physiological reviews, 96 (2016) 911–973.

[2] T. Kusudo, Z. Wang, A. Mizuno, M. Suzuki, H. Yamashita, TRPV4 deficiency increases skeletal muscle metabolic capacity and resistance against diet-induced obesity, Journal of applied physiology, 112 (2012) 1223–1232.

[3] T. Suzuki, T. Notomi, D. Miyajima, F. Mizoguchi, T. Hayata, T. Nakamoto, R. Hanyu, P. Kamolratanakul, A. Mizuno, M. Suzuki, Osteoblastic differentiation enhances expression of TRPV4 that is required for calcium oscillation induced by mechanical force, Bone, 54 (2013) 172–178.

[4] B. Cao, X. Dai, W. Wang, Knockdown of TRPV4 suppresses osteoclast differentiation and osteoporosis by inhibiting autophagy through Ca2+–calcineurin–NFATc1 pathway, Journal of cellular physiology, 234 (2019) 6831–6841.

[5] W. Sun, Y. Luo, F. Zhang, S. Tang, T. Zhu, Involvement of TRP Channels in Adipocyte Thermogenesis: An Update, Frontiers in Cell and Developmental Biology, 9 (2021) 1663.

[6] C.J. O’Conor, T.M. Griffin, W. Liedtke, F. Guilak, Increased susceptibility of Trpv4-deficient mice to obesity and obesity-induced osteoarthritis with very high-fat diet, Annals of the rheumatic diseases, 72 (2013) 300–304.

[7] D.-M. Duan, S. Wu, L.-A. Hsu, M.-S. Teng, J.-F. Lin, Y.-C. Sun, C.-F. Cheng, Y.-L. Ko, Associations between TRPV4 genotypes and body mass index in Taiwanese subjects, Molecular Genetics and Genomics, 290 (2015) 1357–1365.

[8] S. Kumari, A. Kumar, P. Sardar, M. Yadav, R.K. Majhi, A. Kumar, C. Goswami, Influence of membrane cholesterol in the molecular evolution and functional regulation of TRPV4, Biochemical and Biophysical Research Communications, 456 (2015) 312–319.

[9] L.D. Osellame, T.S. Blacker, M.R. Duchen, Cellular and molecular mechanisms of mitochondrial function, Best practice & research Clinical endocrinology & metabolism, 26 (2012) 711–723.

[10] J.H. Lee, A. Park, K.-J. Oh, S.C. Lee, W.K. Kim, K.-H. Bae, The role of adipose tissue mitochondria: regulation of mitochondrial function for the treatment of metabolic diseases, International Journal of Molecular Sciences, 20 (2019) 4924.

[11] R.-h. Lu, H. Ji, Z.-g. Chang, S.-s. Su, G.-s. Yang, Mitochondrial development and the influence of its dysfunction during rat adipocyte differentiation, Molecular biology reports, 37 (2010) 2173–2182.

[12] I. Kladnická, M. Čedíková, M. Kripnerová, J. Dvořáková, M. Kohoutová, Z. Tůma, D. Müllerová, J. Kuncová, Mitochondrial respiration of adipocytes differentiating from human mesenchymal stem cells derived from adipose tissue, Physiological Research, 68 (2019) S287–S296.

[13] D. De Villiers, M. Potgieter, M.A. Ambele, L. Adam, C. Durandt, M.S. Pepper, The role of reactive oxygen species in adipogenic differentiation, Stem cells: Biology and engineering, (2017) 125–144.

[14] E.G. Sustarsic, T. Ma, M.D. Lynes, M. Larsen, I. Karavaeva, J.F. Havelund, C.H. Nielsen, M.P. Jedrychowski, M. Moreno-Torres, M. Lundh, Cardiolipin synthesis in brown and beige fat mitochondria is essential for systemic energy homeostasis, Cell metabolism, 28 (2018) 159–174. e111.

[15] M. Omatsu-Kanbe, K. Inoue, Y. Fujii, T. Yamamoto, T. Isono, N. Fujita, H. Matsuura, Effect of ATP on preadipocyte migration and adipocyte differentiation by activating P2Y receptors in 3T3-L1 cells, Biochemical Journal, 393 (2006) 171–180.

[16] T.K. Acharya, A. Kumar, R.K. Majhi, S. Kumar, R. Chakraborty, A. Tiwari, K.-H. Smalla, X. Liu, Y.-T. Chang, E.D. Gundelfinger, TRPV4 acts as a mitochondrial Ca2+-importer and regulates mitochondrial temperature and metabolism, Mitochondrion, 67 (2022) 38–58.

[17] T.K. Acharya, S. Kumar, T.P. Rokade, Y.-T. Chang, C. Goswami, TRPV4 regulates mitochondrial Ca2+-status and physiology in primary murine T cells based on their immunological state, Life Sciences, (2023) 121493.

[18] T.K. Acharya, A. Kumar, S. Kumar, C. Goswami, TRPV4 interacts with MFN2 and facilitates endoplasmic reticulum-mitochondrial contact points for Ca2+-buffering, Life Sciences, 310 (2022) 121112.

[19] S. Arai, M. Suzuki, S.-J. Park, J.S. Yoo, L. Wang, N.-Y. Kang, H.-H. Ha, Y.-T. Chang, Mitochondria-targeted fluorescent thermometer monitors intracellular temperature gradient, Chemical communications, 51 (2015) 8044–8047.

[20] S. Arai, S.-C. Lee, D. Zhai, M. Suzuki, Y.T. Chang, A molecular fluorescent probe for targeted visualization of temperature at the endoplasmic reticulum, Scientific reports, 4 (2014) 1–6.

[21] X. Liu, T. Yamazaki, H.-Y. Kwon, S. Arai, Y.-T. Chang, A palette of site-specific organelle fluorescent thermometers, Materials Today Bio, 16 (2022) 100405.

[22] T.K. Acharya, S. Kumar, N. Tiwari, A. Ghosh, A. Tiwari, S. Pal, R.K. Majhi, A. Kumar, R. Das, A. Singh, TRPM8 channel inhibitor-encapsulated hydrogel as a tunable surface for bone tissue engineering, Scientific reports, 11 (2021) 1–16.

[23] K. Zebisch, V. Voigt, M. Wabitsch, M. Brandsch, Protocol for effective differentiation of 3T3-L1 cells to adipocytes, Analytical biochemistry, 425 (2012) 88–90.

[24] N.T.S.A.P. ES, A. Miyawaki, Circularly permuted green fluorescent proteins engineered to sense Ca2+, Proc Natl Acad Sci USA, 98 (2001) 3197–3202.

[25] C. Goswami, J. Kuhn, P.A. Heppenstall, T. Hucho, Importance of non-selective cation channel TRPV4 interaction with cytoskeleton and their reciprocal regulations in cultured cells, PloS one, 5 (2010) e11654.

[26] T. Feng, E. Szabo, E. Dziak, M. Opas, Cytoskeletal disassembly and cell rounding promotes adipogenesis from ES cells, Stem Cell Reviews and Reports, 6 (2010) 74–85.

[27] L. Chen, H. Hu, W. Qiu, K. Shi, M. Kassem, Actin depolymerization enhances adipogenic differentiation in human stromal stem cells, Stem Cell Research, 29 (2018) 76–83.

[28] A. Hubry, F.B. Hu, The epidemiology of obesity: A big picture, Pharmacoeconomics, 33 (2015) 673–689.

[29] K. Sarjeant, J.M. Stephens, Adipogenesis, Cold Spring Harbor perspectives in biology, 4 (2012) a008417.

[30] M. Cooper, K. Washburn, The relationships of body temperature to weight gain, feed consumption, and feed utilization in broilers under heat stress, Poultry Science, 77 (1998) 237–242.

[31] A. Güler, H. Lee, I. Shimizu, M. Caterina, Heat-evoked activation of TRPV4 (VR-OAC), J Neurosci, 22 (2002) 6408–6414.

[32] W. Liedtke, D.M. Tobin, C.I. Bargmann, J.M. Friedman, Mammalian TRPV4 (VR-OAC) directs behavioral responses to osmotic and mechanical stimuli in Caenorhabditis elegans, Proceedings of the National Academy of Sciences, 100 (2003) 14531–14536.

[33] L. Landsberg, J.B. Young, W.R. Leonard, R.A. Linsenmeier, F.W. Turek, Do the obese have lower body temperatures? A new look at a forgotten variable in energy balance, Transactions of the American Clinical and Climatological Association, 120 (2009) 287.

[34] L. Ye, S. Kleiner, J. Wu, R. Sah, R.K. Gupta, A.S. Banks, P. Cohen, M.J. Khandekar, P. Boström, R.J. Mepani, TRPV4 is a regulator of adipose oxidative metabolism, inflammation, and energy homeostasis, Cell, 151 (2012) 96–110.

[35] J.M. Ntambi, M. Miyazaki, J.P. Stoehr, H. Lan, C.M. Kendziorski, B.S. Yandell, Y. Song, P. Cohen, J.M. Friedman, A.D. Attie, Loss of stearoyl–CoA desaturase-1 function protects mice against adiposity, Proceedings of the National Academy of Sciences, 99 (2002) 11482–11486.

[36] M. Bishnoi, K.K. Kondepudi, A. Gupta, A. Karmase, R.K. Boparai, Expression of multiple Transient Receptor Potential channel genes in murine 3T3-L1 cell lines and adipose tissue, Pharmacological Reports, 65 (2013) 751–755.

[37] H. Kikuchi, G. Oguri, Y. Yamamoto, N. Takano, T. Tanaka, M. Takahashi, F. Nakamura, T. Yamasoba, I. Komuro, S. Obi, Thermo-sensitive transient receptor potential vanilloid (TRPV) channels regulate IL-6 expression in mouse adipocytes, Cardiovascular Pharmacology: Open Access, (2015).

[38] A.U. Khan, R. Qu, T. Fan, J. Ouyang, J. Dai, A glance on the role of actin in osteogenic and adipogenic differentiation of mesenchymal stem cells, Stem Cell Research & Therapy, 11 (2020) 1–14.

[39] P. Müller, A. Langenbach, A. Kaminski, J. Rychly, Modulating the actin cytoskeleton affects mechanically induced signal transduction and differentiation in mesenchymal stem cells, PloS one, 8 (2013) e71283.

[40] I. Titushkin, S. Sun, A. Paul, M. Cho, Control of adipogenesis by ezrin, radixin and moesin-dependent biomechanics remodeling, Journal of biomechanics, 46 (2013) 521–526.

[41] D.T.V. Diep, K. Hong, T. Khun, M. Zheng, H.-S. Jun, Y.-B. Kim, K.-H. Chun, Anti-adipogenic effects of KD025 (SLx-2119), a ROCK2-specific inhibitor, in 3T3-L1 cells, Scientific reports, 8 (2018) 1–14.

[42] M. Zhai, D. Yang, W. Yi, W. Sun, Involvement of calcium channels in the regulation of adipogenesis, Adipocyte, 9 (2020) 132–141.

[43] J.C. Bournat, C.W. Brown, Mitochondrial dysfunction in obesity, Current opinion in endocrinology, diabetes, and obesity, 17 (2010) 446.

[44] K.V. Tormos, E. Anso, R.B. Hamanaka, J. Eisenbart, J. Joseph, B. Kalyanaraman, N.S. Chandel, Mitochondrial complex III ROS regulate adipocyte differentiation, Cell metabolism, 14 (2011) 537–544.

[45] W. Wang, Y. Zhang, W. Lu, K. Liu, Mitochondrial reactive oxygen species regulate adipocyte differentiation of mesenchymal stem cells in hematopoietic stress induced by arabinosylcytosine, PloS one, 10 (2015) e0120629.

[46] Y. Chen, G.H. Cai, B. Xia, X. Wang, C.C. Zhang, B.C. Xie, X.C. Shi, H. Liu, J.F. Lu, R.X. Zhang, Mitochondrial aconitase controls adipogenesis through mediation of cellular ATP production, The FASEB Journal, 34 (2020) 6688–6702.

[47] Y. Zhang, G. Marsboom, P.T. Toth, J. Rehman, Mitochondrial respiration regulates adipogenic differentiation of human mesenchymal stem cells, PloS one, 8 (2013) e77077.

[48] A. De Pauw, S. Tejerina, M. Raes, J. Keijer, T. Arnould, Mitochondrial (dys) function in adipocyte (de) differentiation and systemic metabolic alterations, The American journal of pathology, 175 (2009) 927–939.

[49] G. Paradies, V. Paradies, V. De Benedictis, F.M. Ruggiero, G. Petrosillo, Functional role of cardiolipin in mitochondrial bioenergetics, Biochimica et Biophysica Acta (BBA)-Bioenergetics, 1837 (2014) 408–417.

[50] A. Prola, F. Pilot-Storck, Cardiolipin Alterations during Obesity: Exploring Therapeutic Opportunities, Biology, 11 (2022) 1638.

[51] M.D. Lynes, F. Shamsi, E.G. Sustarsic, L.O. Leiria, C.-H. Wang, S.-C. Su, T.L. Huang, F. Gao, N.R. Narain, E.Y. Chen, Cold-activated lipid dynamics in adipose tissue highlights a role for cardiolipin in thermogenic metabolism, Cell reports, 24 (2018) 781–790.

[52] L.D. Zorova, V.A. Popkov, E.Y. Plotnikov, D.N. Silachev, I.B. Pevzner, S.S. Jankauskas, V.A. Babenko, S.D. Zorov, A.V. Balakireva, M. Juhaszova, Mitochondrial membrane potential, Analytical biochemistry, 552 (2018) 50–59.

[53] Z. El-Gammal, M.A. Nasr, A.O. Elmehrath, R.A. Salah, S.M. Saad, N. El-Badri, Regulation of mitochondrial temperature in health and disease, Pflügers Archiv-European Journal of Physiology, (2022) 1–9.

[54] D. Chretien, P. Bénit, H.-H. Ha, S. Keipert, R. El-Khoury, Y.-T. Chang, M. Jastroch, H.T. Jacobs, P. Rustin, M. Rak, Mitochondria are physiologically maintained at close to 50 C, PLoS biology, 16 (2018) e2003992.

[55] X. Ma, H. Qian, A. Chen, H.-M. Ni, W.-X. Ding, Perspectives on mitochondria–ER and mitochondria–lipid droplet contact in hepatocytes and hepatic lipid metabolism, Cells, 10 (2021) 2273.

[56] S. Sun, D. Adyshev, S. Dudek, A. Paul, A. McColloch, M. Cho, Cholesterol-dependent modulation of stem cell biomechanics: application to adipogenesis, Journal of biomechanical engineering, 141 (2019).

[57] R. Das, C. Goswami, TRPV4 expresses in bone cell lineages and TRPV4-R616Q mutant causing Brachyolmia in human reveals “loss-of-interaction” with cholesterol, Biochemical and Biophysical Research Communications, 517 (2019) 566–574.

[58] M. Lakk, G.F. Hoffmann, A. Gorusupudi, E. Enyong, A. Lin, P.S. Bernstein, T. Toft-Bertelsen, N. MacAulay, M.H. Elliott, D. Križaj, Membrane cholesterol regulates TRPV4 function, cytoskeletal expression, and the cellular response to tension, Journal of Lipid Research, 62 (2021).

[59] R. Das, C. Goswami, Role of TRPV4 in skeletal function and its mutant-mediated skeletal disorders, Current Topics in Membranes, 89 (2022) 221–246.

[60] R. Das, A. Kumar, R. Dalai, C. Goswami, Cytochrome C interacts with the pathogenic mutational hotspot region of TRPV4 and forms complexes that differ in mutation and metal ion-sensitive manner, Biochemical and Biophysical Research Communications, 611 (2022) 172–178.

[61] J.L. Nourse, V.M. Leung, H. Abuwarda, E.L. Evans, E. Izquierdo-Ortiz, A.T. Ly, N. Truong, S. Smith, H. Bhavsar, G. Bertaccini, Piezo1 regulates cholesterol biosynthesis to influence neural stem cell fate during brain development, Journal of General Physiology, 154 (2022).

[62] A.M. la Rose, V. Bazioti, M. Westerterp, Adipocyte membrane cholesterol regulates obesity, Am Heart Assoc, 2018, pp. 687–689.

[63] E.J. Fernández-Pérez, F.J. Sepúlveda, C. Peters, D. Bascuñán, N.O. Riffo-Lepe, J. González-Sanmiguel, S.A. Sánchez, R.W. Peoples, B. Vicente, L.G. Aguayo, Effect of cholesterol on membrane fluidity and association of Aβ oligomers and subsequent neuronal damage: A Double-Edged Sword, Frontiers in aging neuroscience, 10 (2018) 226.

[64] C. Zhao, Q. Sun, L. Tang, Y. Cao, J.L. Nourse, M.M. Pathak, X. Lu, Q. Yang, Mechanosensitive ion channel Piezo1 regulates diet-induced adipose inflammation and systemic insulin resistance, Frontiers in endocrinology, 10 (2019) 373.

[65] M. Kenmochi, S. Kawarasaki, S. Takizawa, K. Okamura, T. Goto, K. Uchida, Involvement of mechano-sensitive Piezo1 channel in the differentiation of brown adipocytes, The Journal of Physiological Sciences, 72 (2022) 1–15.

[66] C.J. Rendon, E. Flood, J.M. Thompson, M. Chirivi, S.W. Watts, G.A. Contreras, PIEZO1 mechanoreceptor activation reduces adipogenesis in perivascular adipose tissue preadipocytes, Frontiers in Endocrinology, 13 (2022).

[67] S. Wang, S. Cao, M. Arhatte, D. Li, Y. Shi, S. Kurz, J. Hu, L. Wang, J. Shao, A. Atzberger, Adipocyte Piezo1 mediates obesogenic adipogenesis through the FGF1/FGFR1 signaling pathway in mice, Nature communications, 11 (2020) 1–13.

[68] N. Shoham, P. Girshovitz, R. Katzengold, N.T. Shaked, D. Benayahu, A. Gefen, Adipocyte stiffness increases with accumulation of lipid droplets, Biophysical journal, 106 (2014) 1421–1431.

[69] S. Abuhattum, P. Kotzbeck, R. Schlüßler, A. Harger, A. Ariza de Schellenberger, K. Kim, J.-C. Escolano, T. Müller, J. Braun, M. Wabitsch, Adipose cells and tissues soften with lipid accumulation while in diabetes adipose tissue stiffens, Scientific Reports, 12 (2022) 1–17.

[70] B. Glancy, Y. Kim, P. Katti, T.B. Willingham, The functional impact of mitochondrial structure across subcellular scales, Frontiers in physiology, 11 (2020) 541040.

[71] L. Cui, P. Liu, Two types of contact between lipid droplets and mitochondria, Frontiers in cell and developmental biology, 8 (2020) 618322.

[72] M.A. Tarnopolsky, C.D. Rennie, H.A. Robertshaw, S.N. Fedak-Tarnopolsky, M.C. Devries, M.J. Hamadeh, Influence of endurance exercise training and sex on intramyocellular lipid and mitochondrial ultrastructure, substrate use, and mitochondrial enzyme activity, American Journal of Physiology-Regulatory, Integrative and Comparative Physiology, 292 (2007) R1271–R1278.

[73] I.Y. Benador, M. Veliova, K. Mahdaviani, A. Petcherski, J.D. Wikstrom, E.A. Assali, R. Acín-Pérez, M. Shum, M.F. Oliveira, S. Cinti, Mitochondria bound to lipid droplets have unique bioenergetics, composition, and dynamics that support lipid droplet expansion, Cell metabolism, 27 (2018) 869–885. e866.

